# Selective inhibition of human translation by a drug-like compound that traps terminated protein nascent chains on the ribosome

**DOI:** 10.1101/2020.05.20.106807

**Authors:** Wenfei Li, Stacey Tsai-Lan Chang, Fred. R Ward, Jamie H. D. Cate

**Author notes:** Materials & Correspondence. Correspondence and material requests should be addressed to J.H.D.C.

## Abstract

Methods to directly inhibit gene expression using small molecules hold promise for the development of new therapeutics targeting proteins that have evaded previous attempts at drug discovery. Among these, small molecules including the drug-like compound PF-06446846 (PF846) selectively inhibit the synthesis of specific proteins, by stalling translation elongation ^1–4^. These molecules also inhibit translation termination ^4^ by an unknown mechanism. Using cryo-electron microscopy (cryo-EM) and biochemical approaches, we show that PF846 arrests translation at the stop codon by slowing hydrolysis of the protein nascent chain (NC) from peptidyl-site (P-site) tRNA by eukaryotic release factor 1 (eRF1). After NC hydrolysis from the P-site tRNA, PF846 traps the NC in the ribosome exit tunnel in a compact α-helical conformation that induces 28S rRNA nucleotide rearrangements propagating back to the ribosome peptidyl transferase center (PTC). Mutational analyses and human cell-based experiments elucidate the pivotal amino acids of the NC required for PF846-dependent termination arrest, all of which face the PF846 side of the ribosome exit tunnel. The structural and functional data support a model in which PF846 inhibits translation termination by inducing allosteric conformational rearrangements in the NC and PTC that suppress peptidyl-tRNA hydrolysis promoted by eRF1, and trap the NC in the ribosome exit tunnel. This unprecedented mechanism of action reveals new principles of translation termination and lays the foundation for new therapeutic strategies.

## Main

Many diseases are affected by proteins which have been difficult or impossible to target directly with small molecule therapeutics. To treat these diseases, efforts to expand the druggable proteome have attempted to harness knowledge of endogenous cellular pathways to alter gene expression. For example, compounds termed PROteolysis TArgeting Chimeras (PROTACs) serve as molecular recruiters that can direct specific proteins to the ubiquitin-proteasome protein degradation machinery ^5^. Other strategies use small molecules that bind messenger RNAs (mRNAs) to inhibit their translation, or lead to RNA degradation by cellular quality control systems ^6^. Recently, small molecules have even been developed to direct secreted proteins to the lysosome for degradation ^7^. However, ensuring the selectivity of these new small molecule-based strategies is complicated by the fact that they tap into endogenous regulatory pathways with a large number of physiological substrates, requiring a deep understanding of the molecular basis for targeting specificity.

Recently we described the drug-like compound PF846 and its derivatives that target proteins by inhibiting the translation of specific mRNAs by the human ribosome ^1–4^. These small molecules bind in the ribosome exit tunnel in a eukaryotic-specific pocket formed by highly conserved 28S ribosomal RNA (rRNA) residues ^1–4^. Remarkably, they are able to inhibit protein synthesis in a highly selective manner dependent on the nascent polypeptide sequence, opening fundamentally new ways to target proteins of therapeutic interest ^1,4^. Initially discovered to stall translation elongation ^1^, PF846 prevents the movement of mRNA and tRNA on the ribosome by disrupting proper binding of the peptidyl-tRNA in the ribosome peptidyl transferase center ^1,4^.

Surprisingly, PF846 can also block translation termination of specific polypeptides ^4^. Translation termination occurs when an mRNA stop codon enters the ribosome A site. In eukaryotes, termination is mediated by eRF1 and eRF3 (eukaryotic release factors 1 and 3), which form a ternary complex with GTP ^8^. Following stop codon recognition by the eRF1 N-terminal domain, eRF1 rearranges into an extended state that inserts a GGQ-motif (Gly-Gly-Gln) into the PTC to release the nascent peptide ^9–11^. After the nascent peptide is released, the ATPase ABCE1 promotes eRF1 dissociation from the ribosome and ribosome recycling ^12,13^. Differences in the mechanisms of translation elongation and termination suggest PF846 may act in a different manner in these two steps. We therefore used cell-based, structural, and biochemical studies to decipher how the drug-like compound PF846 inhibits translation termination on human ribosomes.

### Selective inhibition of translation termination by PF846 in cells

Earlier we had shown that the nascent chain sequence comprising a segment of human CDH1 (Cadherin-1) followed by the amino acids NPN (Asn-Pro-Asn) (CDH1-NPN*, with * being the stop codon) leads to stalling of the ribosome at the stop codon in a cell-free expression system ^4^. Using this stalling peptide, we first tested whether PF846 also selectively inhibits translation termination in cells. We engineered a stable human cell line (HEK293T) expressing a nanoluciferase reporter fused at its C terminus to the CDH1-NPN polypeptide (**Extended Data Fig. 1a**), along with a second cell line replacing the NPN* motif with GCV* (Gly-Cys-Val, CDH1-GCV*) which is not responsive to PF846 *in vitro* ^4^. Treatment of cells with PF846 showed that the compound inhibits the expression of nanoluciferase fused to CDH1-NPN*, with an IC50 of 3 µM, but does not inhibit expression of nanoluciferase fused to CDH1-GCV*, consistent with *in vitro* translation results (**Fig. 1a and Extended Data Fig. 1**) ^4^.

**Fig. 1.**
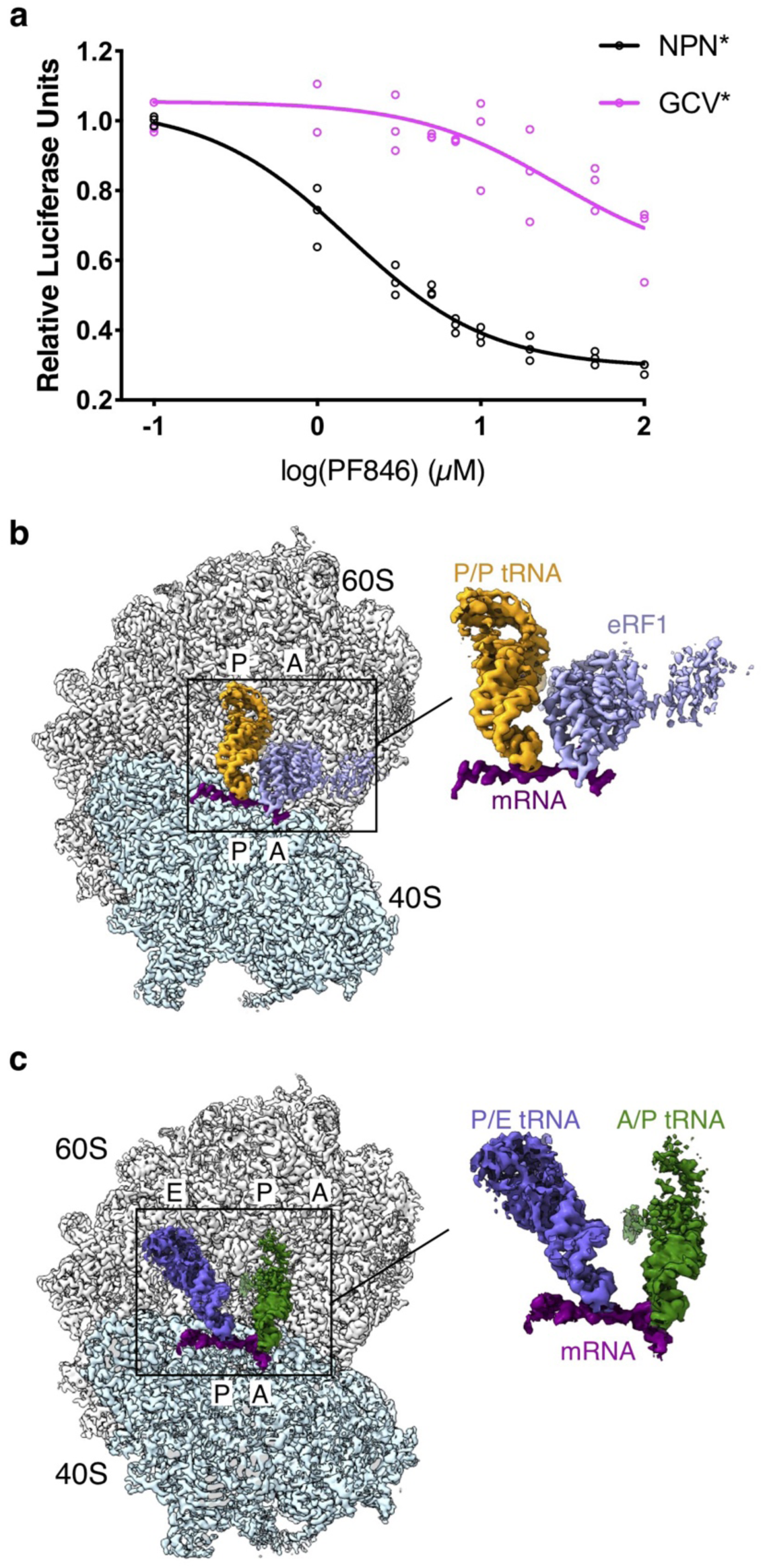
Cell-based assays and structural analysis of PF846-stalled termination complexes. (**a**) Luciferase reporter assays of CDH1–NPN* (black circles) and CDH1– GCV* (magenta circles) stable cell lines in response to different concentrations of PF846 (data present mean ± s.d., *n* = 3 independent experiments). From the CDH1– GCV* data, the IC_50_ of non-specific inhibition is at least 100 μM PF846. (**b**) Cryo-EM reconstruction of CDH1–NPN* RNC in the non-rotated state with the 40S subunit (light cyan), 60S subunit (gray), eRF1 (slate blue) and P/P-site tRNA (orange). A close-up view of the mRNA (magenta), tRNA and eRF1 is shown to the right. (**c**) Structure of CDH1–NPN* RNC in the rotated state bearing A/P-site (dark green) and P/E-site tRNAs (slate blue). A close-up view of the mRNA and tRNAs is shown to the right. Source data for (**a**) is available in **Supplementary Data Set 1**.

### Cryo-EM analysis of a PF846-stalled translation termination complex

To determine the structural basis for how the small molecule PF846 inhibits translation termination, we isolated human ribosome-protein nascent chain (RNC) complexes from *in vitro* translation reactions programmed with an mRNA encoding the CDH1-NPN* fusion protein (**Extended Data Fig. 2a**). We then used cryo-EM to determine structures of the PF846-stalled termination complexes. Particle sorting of the cryo-EM data yielded a major population of ribosomes in the non-rotated state with an overall resolution of 2.8 Å (**Fig. 1b and Extended Data Fig. 2b**), sufficient to visualize chemical modifications of ribosomal RNA (rRNA) and tRNA nucleotides (**Extended Data Fig. 3**) ^14–16^. In the resulting maps, the small molecule PF846 is also well resolved, along with the protein nascent chain, P-site tRNA and eRF1 (**Figs. 1b and 2a, Extended Data Fig. 4**) ^17^. The global conformation of the ribosome resembles that observed previously for eRF1-bound translation termination complexes ^18,19^. A minor population of the RNCs is in the rotated state, bearing tRNAs in the hybrid Aminoacyl site/P site (A/P site) and P site/Exit site (P/E site) (**Fig. 1c, Extended Data Figs. 2b and 5**). The conformation of the RNC complex in the rotated state, including the nascent chain, is reminiscent of that for PF846-stalled RNCs during translation elongation (**Fig. 1c and 2b**) ^4^.

**Fig. 2.**
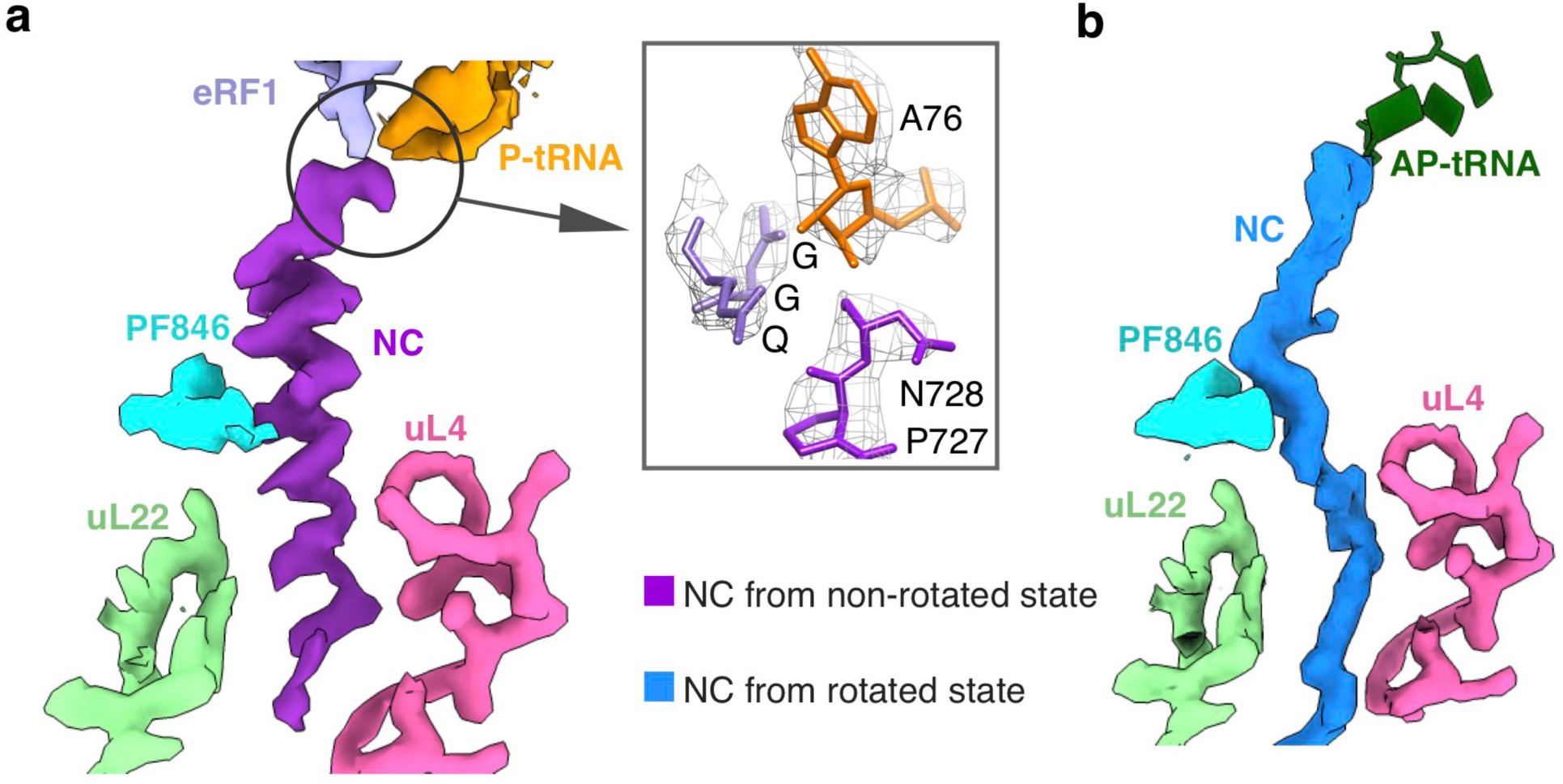
Overview of PF846-stalled nascent chains in the ribosome exit tunnel. (**a**) Cryo-EM density for the stalled nascent chain within the non-rotated RNC (purple). Surface representations of the cryo-EM density for eRF1, P-site tRNA, PF846, and ribosomal proteins are indicated. Inset, view of nascent chain hydrolyzed from P-site tRNA. The model for the GGQ motif of eRF1, A76 of P-site tRNA and the C terminus of the nascent chain is shown in the box with mesh indicating the cryo-EM density. (**b**) The PF846-stalled nascent chain from the rotated RNC (light blue).

### PF846-trapped nascent chain in the ribosome exit tunnel

In the non-rotated state RNC structure, the protein nascent chain is resolved in the ribosome exit tunnel with an average resolution of 3-3.5 Å, allowing us to accurately model the NC conformation and sequence register (**Fig. 2a, Extended Data Fig. 4c**). In striking contrast to the NC in PF846-stalled translation elongation complexes, the NC in the stalled termination complex adopts an α-helical geometry spanning 21 amino acids (residues 705-725), followed by the NPN motif (residues 726-728) in an extended conformation (**Fig. 2a**). Unexpectedly, there is a break in the density between the P-site tRNA and NC, indicating hydrolysis of the ester bond between them (**Fig. 2a**). Consistent with the break in the density, the peptidyl-tRNA is slowly hydrolyzed on the hour timescale in RNCs purified from *in vitro* translation reactions (**Fig. 2a and Extended Data Fig. 6**). These results indicate that the NC remains trapped in the ribosome exit tunnel by PF846 despite the loss of the covalent bond between P-site tRNA and the NC.

In the PF846-stalled RNC in the rotated state, the nascent chain adopts a significantly different conformation from that observed in the non-rotated state. Instead of forming a compact α-helix (**Fig. 2a**), the nascent chain has an extended conformation, as previously seen in PF846-stalled translation elongation complexes (**Fig. 2b**) ^1,4^. Furthermore, the nascent chain density could not be resolved at the amino acid level, as it likely consists of different sequences superimposed on each other ^1,4^. This is supported by the blurred cryo-EM density for the mRNA codon-tRNA anticodon base pairs in the mRNA decoding center (**Extended Data Fig. 7)**, also as previously observed in PF846-stalled translation elongation complexes ^1,4^. Taken together, these structural features are consistent with PF846 slowing translation of the CDH1-NPN polypeptide prior to reaching the stop codon.

### Interactions of the nascent chain with the ribosome exit tunnel and PF846

The unexpected ability of PF846 to trap the NC in the ribosome exit tunnel even after peptidyl-tRNA hydrolysis (**Figs. 2 and 3**) led us to investigate the contribution of single amino acid residues of the NC during PF846-induced stalling. We previously identified the NPN motif (residues 726-728) in an mRNA library-based experiment, in which a range of sequences in these positions supported inhibition of translation termination ^1,4^. Within the library, the amino acid in position 728 could accommodate nearly all other amino acids except for the largest two, tyrosine and tryptophan (Y and W) ^1,4^. In support of these findings, N728 projects towards a pocket formed by 28S rRNA residues G3886-C3888 that leaves room for larger amino acid side chains (**Fig. 3b**). P727 makes multiple contacts to U4422 and Ψ4500 in 28S rRNA (**Fig. 3c**), and cannot tolerate larger amino acids, again consistent with enrichment of predominantly P and V at this position ^1,4^. N726 makes hydrogen bond and van der Waals contacts to U4422 and A3887, and can tolerate replacement with amino acids D and H, as seen in the previously-identified sequence motif (**Fig. 3d**) ^1,4^.

**Fig. 3.**
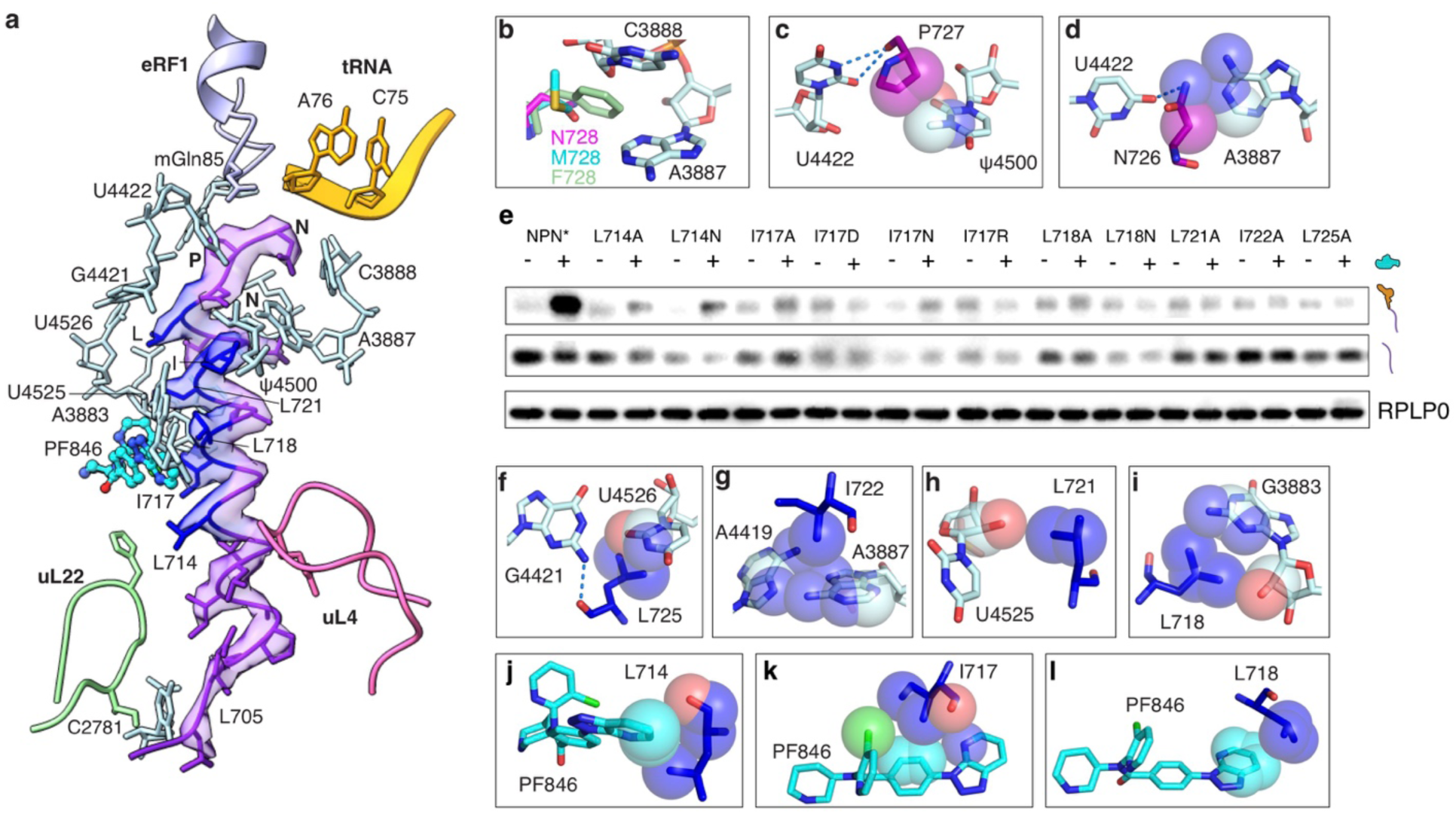
Structural details of the nascent chain in the non-rotated state and effects of mutations in the NC. (**a**) The CDH1-NPN nascent chain model within the cryo-EM density (transparent surface). rRNA bases (light cyan) and ribosomal protein residues (light green for uL22 and hot pink for uL4) that have close interactions with the nascent chain are shown. The amino acids of the nascent chain found to be essential for PF846-mediated stalling are colored blue. Amino acids without numbers are: I722, L725, N726, P727, and N728 at the C-terminus. (**b**) Models of different amino acids at the NC C-terminal position 728. The pocket formed by 28S rRNA nucleotides A3887 and C3888 can accommodate different sizes of amino acid, with methionine (M728) and phenylalanine (F728) as examples. (**c**-**d**) Interactions between P727 and N726 with adjacent 28S rRNA nucleotides. Dashed lines indicate hydrogen bonds and spheres represent van der Waals radii of C, N and O atoms. (**e**) Western blots of FLAG-tagged CDH1-NPN nascent chains containing single mutations, from *in vitro* translation reactions in the presence (+) or absence (–) of 50 μM PF846. The positions of tRNA-bound and free nascent chains are shown, with RPLP0 serving as a loading control. (**f**-**i**) Interactions between 28S rRNA nucleotides and the PF846-stalled nascent chain in the non-rotated RNC with dashed lines indicating hydrogen bonds and spheres representing van der Waals radii. (**j**-**l**) Direct interactions between PF846 and nascent chain residues. Spheres represent van der Waals radii. O, N and Cl atoms are colored in red, blue and green, respectively. Uncropped gel images for (**e**) are available in **Supplementary Data Set 2**.

To test whether the NPN sequence is sufficient for PF846-dependent inhibition of translation termination, we examined the NPN motif in the context of PCSK9 (Proprotein convertase subtilisin/kexin type 9). Notably, the predicted secondary structure of the PCSK9 sequence originally identified as sufficient for PF846 to stall translation elongation ^1,4^ is unlikely to form a compact α-helical geometry as observed in the CDH1-NPN* structure (**Extended Data Fig. 8a**). We inserted the NPN* motif at different positions in the PCSK9 sequence (**Extended Data Fig. 8**). In contrast to the effects of PF846 on PCSK9 during translation elongation, PF846 was unable to inhibit translation of any of the C-terminal PCSK9-NPN* sequences (**Extended Data Fig. 8**).

Consistent with the observation above that the NPN motif is necessary but not sufficient for PF846-dependent inhibition of translation termination, mutation of L725 in the CDH1-NPN NC, abolishes translational stalling (**Fig. 3e**). In the structure of the stalled termination complex, L725 makes a backbone hydrogen bond with G4421 and side chain contacts to U4526 (**Fig. 3a and f**). In the N-terminal direction of the NC preceding L725, one face of the long α-helical segment of the NC makes numerous contacts with nucleotides lining the ribosome exit tunnel and PF846 (**Fig. 3a**). Nascent chain residues L718, L721 and I722 contact 28S rRNA nucleotides U4525, A4419, A3887 and G3883 (**Fig. 3a, g-i**), and residues L714, I717 and L718 engage in multiple interactions with PF846 through van der Waals forces (**Fig. 3j-l**). These interactions are essential for the PF846-induced translation termination as reflected by the mutagenesis results (**Fig. 3e**). In the distal region of the ribosome exit tunnel (i.e. away from the PTC and further towards the NC N-terminus), the NC makes contacts with ribosomal protein uL4 (residues W67 and R71) and uL22 (residues H133 and I136), along with nucleotide C2781 in 28S rRNA (**Fig. 3a and Extended Data Fig. 9a-d**). However, mutational analyses reveal that ribosome contacts to NC residues L705-L711 are less important for the inhibition of translation termination (**Extended Data Fig. 9e-f**).

To further test the importance of contacts of the NC to PF846 and the distal region of the ribosome exit tunnel, we constructed two cell lines expressing nanoluciferase-CDH1-NPN* reporters harboring two representative NC mutations, I717A and Q706A. Whereas the I717A mutation decreased the overall magnitude of PF846-induced stalling and increased the IC50 ∼3x compared to the CDH1-NPN* sequence, the Q706A mutation had no effect on overall inhibition (**Extended Data Fig. 9g**). Taken together, the structural and mutagenesis results reveal that PF846-induced inhibition of translation termination requires a NC sequence distributed over 15 amino acids (residues L714-N728).

### eRF1 conformation in the PF846-stalled termination complex

Although structurally different, translation release factors (RFs) from bacteria to eukaryotes all recognize the nucleotides in stop codons and catalyze peptide release ^20–22^. They possess a highly conserved GGQ sequence motif that resides at the tip of a short α-helix and points directly into the PTC (**Fig. 4a**). The Gln residue is modified to *N*^5^-methyl-Gln (mGln) in both the bacterial and eukaryotic domains ^23^. Computational analysis and recent structural studies of bacterial release factor 2 (RF2) showed that this methylation can enhance the correct positioning of the Gln residue in the PTC catalytic site during translation termination ^11,24^. However, in eukaryotes, the positioning of mGln remains unclear due to the lack of a high resolution structure of a termination complex ^9,10,21,25^. Here, in the PF846-stalled translation termination RNCs, we clearly resolved the density for the mGln of eRF1 (**Extended Data Fig. 10a**). In bacteria, the *N*5-methylation increases the van der Waals interactions with 23S rRNA nucleotides in the PTC, including *Ec* U2506, *Ec* A2451 and *Ec* A2452 (*Ec, E. coli* numbering, **Fig. 4b**) ^11,26^. In PF846 stalled translation termination complex, mGln adopts the same conformation and establishes multiple interactions with the equivalent PTC residues in human 28S rRNA (**Fig. 4a-c and Extended Data Fig. 10a**).

**Fig. 4.**
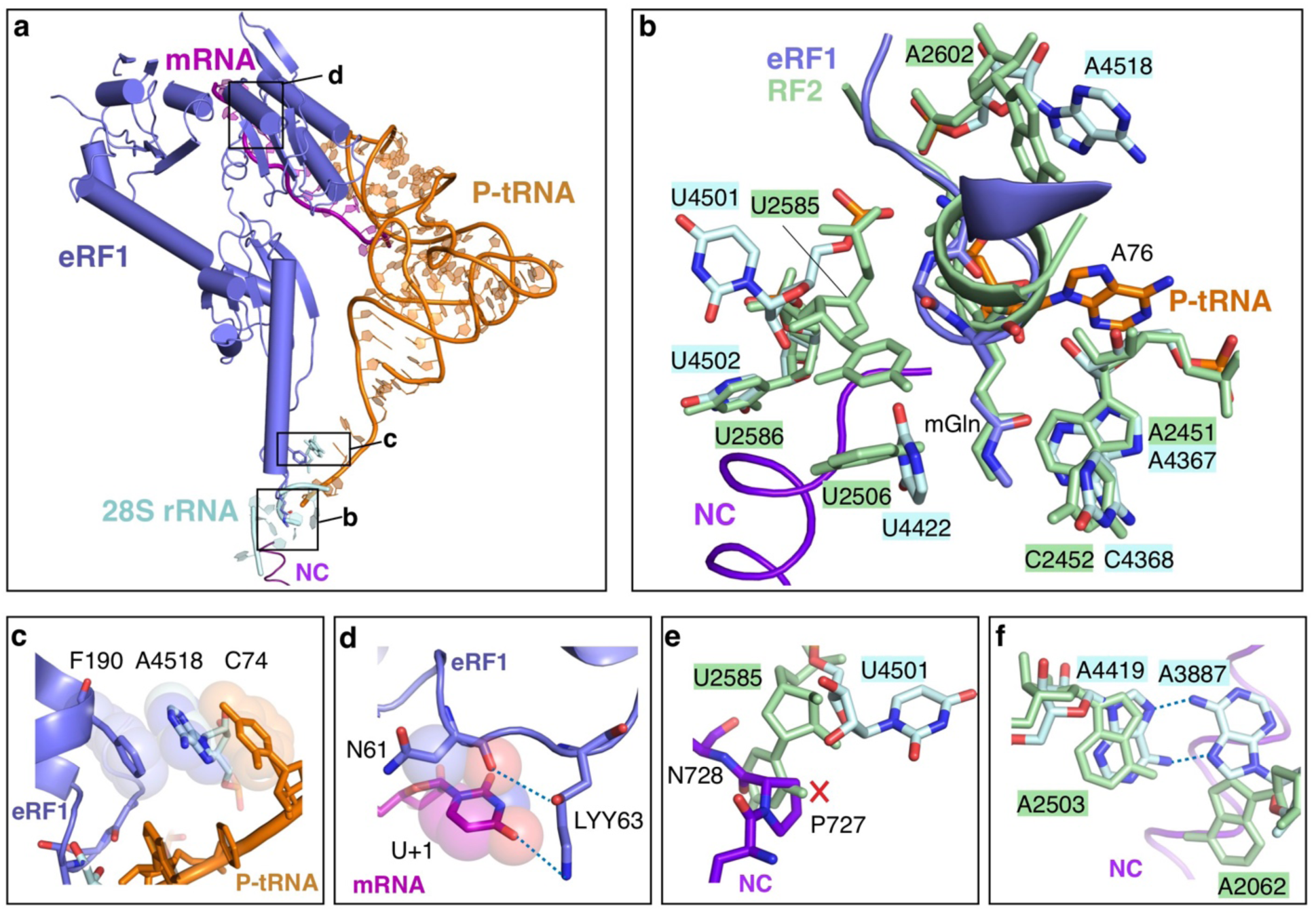
PTC rearrangements in the PF846-stalled termination complex. (**a**) Overview of the mRNA decoding center and PTC with eRF1 (slate blue), P-site tRNA (orange), mRNA (magenta), 28S rRNA (light cyan), nascent chain (purple) and PF846 (cyan). Essential eRF1 motifs are highlighted in boxes with letters corresponding to the panel labels. (**b**) Comparison of the PTC from a bacterial-RF2 ribosome complex (PDB code 6c5l ^32^, pale green) and in the PF846-stalled termination complex. (**c**) eRF1-mediated stacking interactions within the PTC, showing eRF1 residue F190, 28S rRNA nucleotide A4518, and C74 of P-site tRNA. Spheres represent van der Waals radii of C, N and O atoms. (**d**) Interaction network of U+1 in the UAA stop codon (magenta) with the eRF1 NIKS motif (slate blue). Interactions proposed to stabilize the stop codon are indicated with dashed lines for hydrogen bonds and spheres for van der Waals radii. 4*R*-hydroxylysine (LYY63) at residue 63 in eRF1 is shown. (**e**) Comparison of position of 28S rRNA nucleotide U4501 with *Ec* 23S rRNA nucleotide U2585 in the RF2-bound bacterial ribosome. The steric clash with the PF846-stalled NC is highlighted with a red “X”. (**f**) Comparison of positions of A3887 and A4419 with the respective nucleotides in the RF2-bound bacterial ribosome (*Ec* A2062 and U2503), showing dashed lines for hydrogen bonds.

In the present structure, eRF1 docks in the mRNA decoding site, positioning a highly conserved NIKS motif (Asn-Ile-Lys-Ser) located in the N-terminal domain to recognize the stop codon (**Fig. 4a and d**) ^27^. During translation termination, eRF1 is thought to directly interact with the uridine at position 1 in the stop codon (U1) by means of a post-translationally modified lysine (residue K63 in the NIKS motif) hydroxylated at the C4 carbon ^28^. This modification increases translation termination efficiency by reducing stop codon read-through ^29^. Although prior structures modeled K63 within hydrogen bonding distance of U1 ^9,10^ the limited resolution of the cryo-EM density precluded modeling of the 4-hydroxylation. In the present structure, we were able to model 4*R*-hydroxylysine at residue 63 in the cryo-EM density, with the hydroxyl group hydrogen bonding to the backbone carbonyl of asparagine 61 in the NIKS motif and the side chain ε-amine hydrogen bonding to the O4 carbonyl in U1 (**Fig. 4d and Extended Data Fig. 10b**).

### Ribosomal RNA elements critical for PF846-induced inhibition of termination

The positioning of eRF1 in both the mRNA decoding site in the small ribosomal subunit and the PTC in the large ribosomal subunit indicates eRF1 is bound in an active conformation ^9–11^. However, rearrangements in the PTC and ribosome exit tunnel could explain the slow hydrolysis of peptidyl-tRNA and trapping of the nascent chain by PF846. In bacteria two universally conserved rRNA nucleotides, *Ec* A2602 and *Ec* U2585 (A4518 and U4501 in human 28S rRNA, respectively) are required for peptide release ^30,31^. In the present structure, A4518 occupies a similar position to *Ec* A2602 in the bacterial translation termination complex ^32^, and stabilizes the positioning of the GGQ catalytic loop by stacking between F190 in eRF1 and the C74 of P-site tRNA (**Fig. 4c**). However, nucleotide U4501 (*Ec* U2585) is rotated by 90° away from the PTC, to avoid a steric clash with P727 in the nascent chain (**Fig. 4b and e, Extended Data Fig. 11b**). Interestingly, the same change in the position of U4501 is observed both in human cytomegalovirus (hCMV) arrested translation termination complexes ^10^ (**Fig. 5a, b and Extended Data Fig. 11b**) and macrolide-dependent stalling of ErmCL in bacteria ^33,34^ In the ribosome exit tunnel, universally conserved nucleotides A3887 and A4419 (*Ec* A2062 and *Ec* A2503, respectively) form a non-canonical A-A base pair (**Fig. 4f**) previously found to be essential in macrolide-dependent ribosome stalling (**Fig. 5c**) ^30,33^. In this conformation, A3887 and A4419 make multiple interactions with NC residues essential for PF846-induced inhibition of termination (**Fig. 3b, d and g**). Reflecting the sequence specificity of stalling, A3887 adopts a conformation that avoids a steric clash with the NC, distinctly different from that observed in hCMV-induced stalling (**Extended Data Fig. 11c**). Finally, U4422 (*Ec* U2506), which is known to move positions as part of the induced fit of the PTC required for peptide bond formation ^35,36^, is highly mobile based on the cryo-EM density, forming different extents of contacts to the nascent chain and mGln185 of eRF1 (**Extended Data Fig. 11d-h**). This suggests U4422 may be less essential in PF846-mediated trapping of the CDH1-NPN NC on the ribosome.

**Fig. 5.**
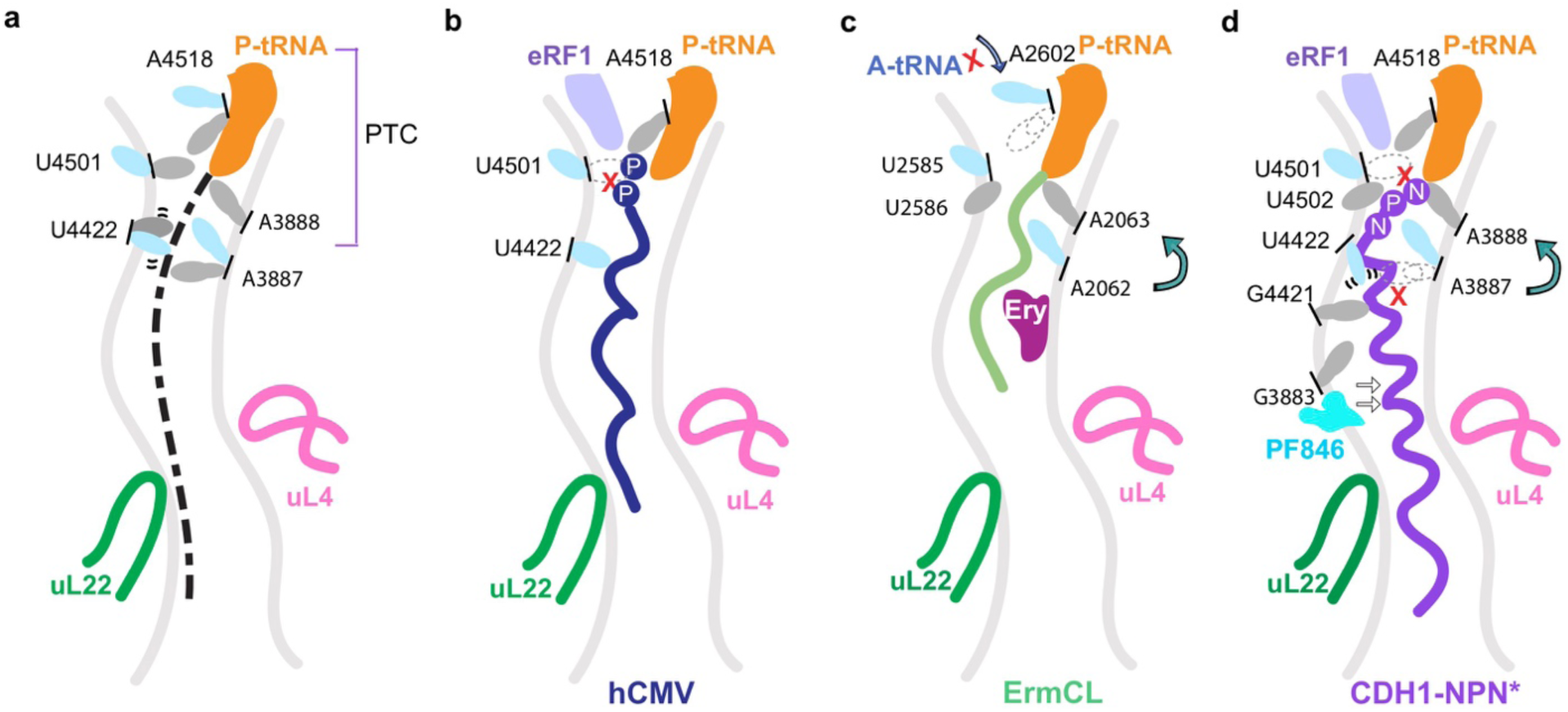
Features of PF846-mediated inhibition of translation termination. (**a**) Schematic of common translation stalling features, with 28S rRNA nucleotides showing conformational changes (grey, translation-competent conformation; light cyan, inactive conformation). Nucleotides are shown with human numbering, with the corresponding *E. coli* numbering in parentheses as follows: A4518 (*Ec* A2602), U4501 (*Ec* U2585), U4422 (*Ec* U2506), A3888 (*Ec* A2063), A3887 (*Ec* A2062). (**b**) Schematic showing hCMV nascent-chain induced stalling of translation termination ^10^. The critical amino acid P-P motif for hCMV-mediated stalling is indicated. U4501 is flipped away from the PTC to avoid a steric clash with the C-terminal motif, leading to inhibition of PTC catalysis. (**c**) Schematic representing drug-dependent stalling of ErmCL in bacteria ^30^. A2602 blocks the accommodation of A-site tRNA, and U2585 flips away, inactivating the PTC. A2062 in the ribosome exit tunnel exhibits a distinct conformation due to the geometry of the NC and transmits the stalling signal back to the PTC through the neighboring nucleotides including A2063. (**d**) Model for PF846-stalled translation termination. The ribosome exit tunnel nucleotides essential for stalling are labeled. Stalling is propagated from PF846 back to the PTC by contacts with and rearrangements of multiple nucleotides. In panels **b**-**d**, collisions with nucleotides in their active conformation are shown by a red “X”.

## Discussion

We recently reported that drug-like compounds can selectively stall translation during nascent peptide elongation, by impeding the movement of the ribosomes along the mRNA in a nascent chain-dependent manner ^1,4^. These compounds exert their selective effect while bound in the ribosome exit tunnel, despite the fact that all protein nascent chains transit this space in the ribosome. Unexpectedly, these same small molecules can also block eukaryotic translation termination on specific nascent chain sequences ^1,4^. Here we show these molecules employ a unique mechanism of action to block translation termination and trap the released protein nascent chain on the ribosome.

Using cryo-EM we identified two different states of the stalled RNCs, the first in a rotated state similar to that observed for PF846-stalled translation elongation (**Figs. 1c and 2b**) ^4^. The second state, a post-hydrolysis complex, contains the nascent chain hydrolyzed from the P-site tRNA yet trapped in the ribosome exit tunnel (**Fig. 1b, Fig. 2a**). These two structures suggest a model for PF846-mediated inhibition of translation termination. First, PF846 induces a slowdown in translation before the stop codon enters the mRNA decoding site (A site), similar to what is observed for PF846-mediated stalling of translation elongation ^4^ (**Fig.1c and Extended Data Fig. 7d-f**). Subsequently, after the stop codon is recognized by eRF1, the NC is slowly hydrolyzed from the P-site tRNA but is not released from the RNC (**Fig. 2a and Extended Data Fig. 6**). The NC adopts a highly compact α-helix conformation within the ribosome exit tunnel, and remains trapped in the RNC due to multiple interactions with the ribosomal exit tunnel wall and PF846 (**Fig. 3**). Notably, the ability of PF846 to inhibit translation termination in cells (**Fig. 1a and Extended Data Fig. 1**) suggests its interactions with the compact nascent chain traps the nascent chain in the ribosome despite pulling forces that may be exerted by chaperones or cotranslational protein folding (**Fig. 2a, Fig. 3 and Extended Data Fig. 6**) ^19,37^.

Stalling of translation termination by PF846 involves mechanisms distinct from those seen in bacterial and other eukaryotic stalling systems (**Fig. 4 and Fig. 5**) ^10,38–41^. For example, the hCMV stalling peptide forms an α-helix in the exit tunnel, but translation termination is mainly inhibited by two C-terminal prolines (**Fig. 5b**) ^9,10,42^. By contrast, the C-terminal NPN motif by itself is insufficient for PF846-induced stalling (**Fig. 3**). Instead, PF846 traps the NC in the ribosome exit tunnel by making contacts along one face of a longer NC α-helix (**Fig. 5d and Extended Data Fig. 12**). Mutation of these residues leads to diminished or abolished stalling (**Fig. 3**). Additional contacts between the NC and nucleotide A3887 (*Ec* A2062) in the exit tunnel are required for PF846-induced stalling (**Fig. 3**), similar to macrolide-dependent stalling in bacteria (**Fig. 5c-d**) ^30^. However, in contrast with macrolide-dependent stalling in bacteria, PF846 does not lead to nucleotide rearrangements that block access to the A site in the PTC (**Fig. 5c-d**). The distributed sequence determinants important for PF846-induced stalling of translation termination (8 of the C-terminal NC amino acids within the ribosome exit tunnel) indicate that molecules like PF846 could be used for selective inhibition of translation for new therapeutic targets. Taken together, our experimental data elucidate the mechanism of drug-like compound stalled translation termination in human ribosomes, providing insights into the sequence specificity of compounds like PF846 that can aid development of these compounds for therapeutic purposes.

## Supporting information

Supplementary Data Set 1

Supplementary Data Sets 2-5

## Acknowledgements

We thank D. Toso (Bay Area Cryo-EM consortium) and P. Tobias for help with microscope operation and data collection; D. De Silva, M. Pulos-Holmes and Z. Zhang in sharing the lentivirus related plasmids and suggestions in making the stable cell lines; Z. Watson for helpful discussions in Cryo-EM data processing; N. Aleksashin for valuable comments on the manuscript. This work was funded by the NIH (grants R01-GM065050 and R01-GM131142).

## Contributions

W.L. performed the sample preparation, acquired cryo-EM data, carried out image processing and structure refinement. W.L. and J.H.D.C. did the model building and structural analysis of the modified rRNA and amino acids. W.L. and S.T.-L.C. conducted the biochemistry and cell-based experiments. W.L. and J.H.D.C. wrote the original draft of the manuscript. All authors analyzed the data and edited the manuscript.

## Methods

### DNA constructs and *in vitro* transcription

The DNA plasmid encoding the mRNA for the CDH1-NPN* stalling construct was previously described ^4^, which includes CDH1 residues 586-725, followed by amino acids NPN and a UAA stop codon (**Extended Data Fig. 2a**). Point mutations in the CDH1-NPN* protein nascent chain sequences were generated through “around-the-horn” cloning ^43^ using primers with overhangs (**Extended Data Table 1**). Plasmids used to construct lentiviral vectors encoded nanoluciferase with an N-terminal 3XFLAG peptide, and the CDH1 stalling sequence with different termination motifs or mutations (GCV*, NPN*, Q706A and I717A), along with the CDH1 3’-untranslated region (3’-UTR) (**Extended Data Fig. 1a**). The DNA fragment was assembled using overlap extension PCR with primers shown in **Extended Data Table 1**. The resulting PCR product was subsequently inserted using Gibson Assembly Master Mix (NEB, E2611L) into the lentiviral vector CD813A (System Biosciences) encoding the 5’ untranslated region of human β-globin (*HBB* 5’-UTR).

DNA templates for *in vitro* transcription were amplified by PCR using primers encoding a T7 RNA polymerase promoter and a poly-A tail (**Extended Data Table 1**). All PCR products were purified via QIAquick Gel Extraction Kit (Qiagen, 28115) before their use in *in vitro* transcription reactions. Messenger RNAs were transcribed using T7 RNA polymerase prepared in house. Reactions were set up with 20 mM Tris-HCl pH 7.5, 35 mM MgCl_2_, 2 mM spermidine, 10 mM DTT, 1 U/mL inorganic pyrophosphatase (ThermoFisher, EF0221), 7.5 mM each NTP, 0.2 U/L SUPERaseIn RNase Inhibitor (ThermoFisher, AM2696), 0.1 mg/mL T7 RNA polymerase and 40 ng/μL DNA. After 4 hours incubation at 37 °C, 0.1 U/μL RQ1 RNase-free DNase (Promega, M6101) was added to the reactions, followed by another incubation at 37 °C for 30 min to remove the template DNA. RNA was precipitated overnight at -20 °C after adding 1/2 volume of 7.5 M LiCl/50 mM EDTA, and the resulting pellet was washed with cold 70% ethanol and dissolved with RNase free water. RNAs were purified using Zymo RNA Clean and Concentrator kit following the manufacturer’s instructions (Zymo research, R1017) before use in *in vitro* translation reactions.

### Generation of stable cell lines

Lentiviruses encoding the various CDH1-derived stalling sequences were generated using HEK293T cells in 10 cm dishes. Cells grown to a confluence of 80% were transfected with the CD831A plasmids encoding the stalling sequences described above, together with helper plasmids PsPAX2 and pCMV-VSV-G (Addgene), using the TransIT-LT1 transfection reagent (Mirus Bio, MIR 6000) following the manufacturer’s instructions. Lentiviruses were harvested and filtered with 0.22 μm filters after 48 hours and 72 hours. To generate stable cell lines, 1 mL of each virus was added to HEK293T cells seeded in a 6-well plate at a density of around 80%-90% confluence, along with 10 μg/μL of polybrene (Millipore, TR-1003-G). After 24 hours, the cells were treated with 4 μg/mL puromycin (Gibco, A1113803) for 4 days and split in DMEM media (Dulbecco’s Modified Eagle Medium, Invitrogen) with 10% FBS (Tissue Culture Biologicals) before use.

### Luciferase reporter assay

To determine the steady-state conditions for PF846-induced stalling of nanoluciferase reporters, the stable cell lines expressing nanoluciferase reporter genes fused to the various CDH1 stalling sequences were treated with 5 μM PF846 in 0.1% DMSO or 0.1% DMSO as the control. Luciferase activity was assayed after 1.5, 7, 10, 20, 27 and 43 hr using the Nano-Glo® Luciferase Assay System (Promega, N1120). To determine the concentration dependence of PF846 inhibition of nanoluciferase reporters, the cell lines were treated with 0.1% DMSO and different concentrations of PF846 in 0.1% DMSO (1, 3, 5, 7, 10, 20, 50 and 100 μM) for 20 hr. After treatment, the luciferase activity was assayed as described above. The IC_50_ calculated here is likely an upper bound due to the accumulation of secreted nanoluciferase in the cell culture media ^44^.

### *In vitro* translation reactions

Extracts from HeLa cells were made as described previously ^1,4^. For the *in vitro* translation reactions to assess mutations in the nascent chain sequence, a 30 μL final volume was used for each mRNA, which contained 15 μL cell lysate and buffer with a final concentration of 20 mM HEPES pH 7.4, 120 mM KOAc, 2.5 mM Mg(OAc)_2_, 1 mM ATP/GTP, 2 mM creatine phosphate (Roche), 10 ng μL−1 creatine kinase (Roche), 0.21 mM spermidine, 0.6 mM putrescine, 2 mM TCEP (tris(2-carboxyethyl)phosphine), 10 μM amino acids mixture (Promega, L4461), 1 U μL−1 murine RNase inhibitor (NEB, M0314L), 600 ng mRNA and 50 μM PF846 in 1% DMSO, or 1% DMSO as control. Translation reactions were incubated for 23 min at 30 °C, followed by centrifugation for 5 min at 20,000 *g* to remove the cell debris. The supernatant was applied to a 50% sucrose cushion prepared with cushion buffer (25 mM HEPES-KOH pH 7.5, 120 mM KOAc and 2.5 mM Mg(OAc)_2_, 1 M sucrose, 1 mM DTT, 50 μM PF846) and centrifuged for 1 hour at 603,000 *g* using a MLA-130 rotor (Beckman Coulter) at 4 °C. The pellet was suspended in ice-cold RNC buffer (20 mM HEPES pH 7.4, 300 mM potassium acetate, 5 mM magnesium acetate, 1 mM DTT, 0.2 mM PF846). The RNC samples were then used for western blot analysis.

### RNase A treatment and western blot

To assess the total amount of stalled nascent chains, RNC samples purified from sucrose cushions as described above were treated with 100 μg mL^-1^ DNase-free RNase A (ThermoFisher, EN0531) at 37 °C for 50 min followed by western blot with monoclonal Anti-FLAG M2-Peroxidase (HRP) antibody (Sigma, A8592) for the CDH1-derived nascent chains (1:10,000 dilution), as well as with an anti-RPLP0 antibody (Bethyl Laboratories, A302-882A) used as a loading control (1:8,000 dilution). In order to visualize tRNA-bound stalled nascent chains, the RNC samples from the above sucrose cushions were heated at 55 °C for 5 min in the presence of 1X Laemmli Sample buffer (Bio-Rad, 1610737). Subsequently, these samples were resolved on NuPAGE 4-12% Bis-Tris protein gels (Thermo Fisher Scientific, NP0323PK2) prior to western blot analysis.

### Purification of stalled RNCs and Cryo-EM grid preparation

An *in vitro* translation reaction of 1.5 mL programmed with mRNA encoding the CDH1-NPN* sequence was incubated with 50 μM PF846 at 30 °C for 23 min and then centrifuged at 20,000 *g* for 5 min. The supernatant was incubated with 50 μL of anti-FLAG M2 agarose beads (Sigma, A2220) for 1 hour at room temperature with gentle mixing. All the following purification steps were conducted at room temperature unless specifically noted. The 3XFLAG-tagged RNCs bound to the anti-FLAG beads were washed 3 times with 200 μL RNC buffer, then 3 times with 200 μL RNC wash buffer plus 0.1% TritonX-100, followed by 3 times with 200 μL RNC buffer plus 0.5% TritonX-100, and finally washed twice with 200 μL RNC buffer. A final concentration of 0.2 mg mL^-1^ 3X FLAG peptide (Sigma, F4799) in the RNC buffer was used to release the RNCs from the beads. The resulting RNCs were pelleted through a sucrose cushion as described above, and resuspended in ice-cold RNC buffer which were immediately used for making cryo-EM grids.

The concentration of the purified RNC used for cryo-EM grid preparation was 50 nM. Approximately 3.2 μL of RNCs were incubated on plasma-cleaned 300-mesh holey carbon grids (C-flat R2/2, Electron Microscopy Science) for 1 min, on which a home-made continuous carbon film was pre-coated. Grids were blotted for 3 sec under 10% humidity at 4 °C and plunge-frozen in liquid ethane using a FEI Vitrobot.

### Cryo-EM data collection and processing

Cryo-EM data were collected using a Titan Krios electron microscope (FEI) equipped with a K2 Summit direct detector and GIF Quantum filter (Gatan) at 300 kV (**Table 1**). The total exposure time was 9 sec, with a total dose of 50 electrons Å^-2^ (frame dose 1.3 electrons Å^-2^). The frames in the resulting movies were corrected for motions using MotionCor2 with FtBin 2 and dose weighting ^45^. The subsequent processing was performed in RELION 3.1 ^46^. For the initial steps of image processing, the data were binned by a factor of 4. After 2D classifications to remove images with ice or other contaminants, 3D classification was used to remove non-ribosomal particles (82,787 particles). After 3D refinement, an additional 3D classification was performed to separate the ribosomes in the rotated state (26,463 particles) and non-rotated state (57,324 particles) (**Extended Data Fig. 2b**). After one round of CTF refinement and Bayesian polishing ^47^, particles of each class were further processed by focused refinements with individual masks applied (**Extended Data Fig. 2b**) ^48^. Binary masks with smoothed edges were generated with the “relion_mask_create” tool in RELION. Post-processing and B-factor sharpening implemented in RELION3.1 was applied to the final maps. EMBFACTOR was also used to apply certain B-factors in order to better visualize the density of certain parts of the RNC maps ^49^. The reported resolutions for all maps is based on the FSC cutoff criterion of 0.143 ^50,51^. Local resolution estimation was performed using Resmap ^52^.

**Table 1.**
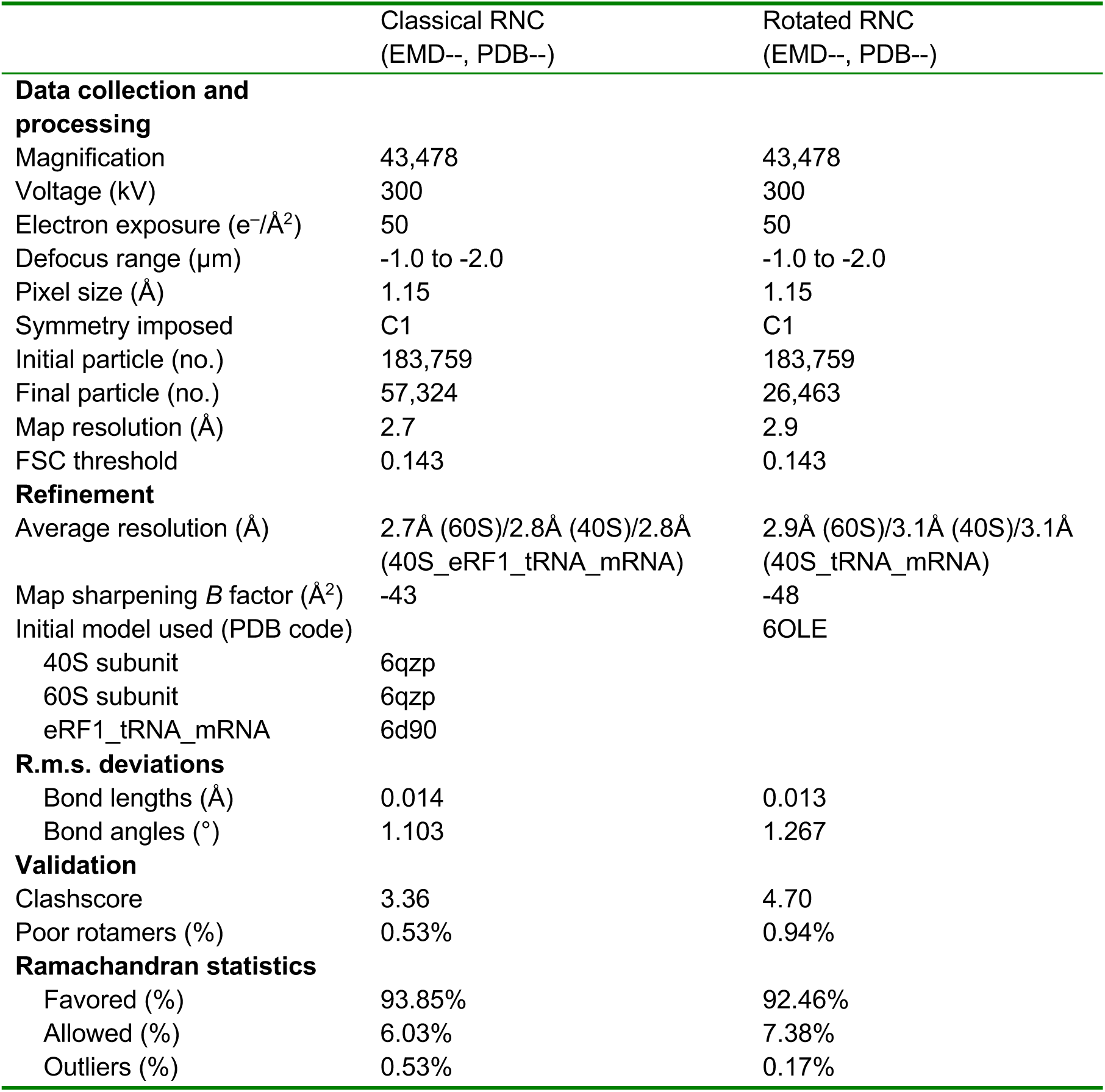
Cryo-EM data collection, refinement and validation statistics of PF846-stalled termination complexes.

### Model building and refinement

The atomic models of the human 80S ribosome (PDB: 6qzp) ^14^ were used as the starting point and manually adjusted and refined in COOT ^53^ using the experimental cryo-EM maps. For the non-rotated RNC, eRF1 and P-tRNA was initially modeled using the mammalian 80S ribosome with eRF1 and P-site tRNA bound (PDB: 6D90) ^54^. The CDH1-NPN nascent chain was modeled manually into the density using COOT ^53^ (CDH1-NPN nascent chain sequence: EAGLQIPAILGILGGILALLILILNPN). The post-translational modification of K63 in eRF1, 4-hydroxylysine (LYY63), was generated using ChemDraw 19.0 (PerkinElmer) to create the SMILES string, which was then used as the input to phenix.elbow ^55^. Subsequent adjustments to the eRF1 model in the vicinity of the 4-hydroxylysine were made using phenix.real_space_refine ^55–57^ and the 40S subunit focused-refined map. The post-translational modification of Q185 to mGln185 used the standard .cif library in CCP4 ^58^.

To model the rotated-state RNC structure, we used the previously-reported PF846-stalled RNC model (PDB: 6OLE) ^4^ docked into the cryo-EM map by rigid-body fitting followed by refinement in Phenix ^56,57^. Both RNC structures were refined using Phenix (phenix.real_space_refine) with RNA secondary structure restraints imposed ^56,57^. Model refinement and validation statistics are provided in **Table 1**.

### Figure preparation

Figures were prepared using UCSF Chimera ^59^, UCSF ChimeraX ^60^, and PyMOL (Schrödinger).

### Secondary structure modeling

Modeling of the secondary structure of the PF846-sensitive PCSK9 stalling sequence was carried out using Phyre2 ^61^.

### Reporting Summary

Further information on research design is available in the Nature Research Reporting Summary linked to this article.

### Data availability

The cryo-EM maps have been deposited with the Electron Microscopy Data Bank under the accession codes EMD-____(non-rotated RNC with eRF1) and EMD-____(rotated RNC). Atomic Coordinates have been deposited in the Protein Data Bank with accession codes____(non-rotated RNC with eRF1) and____(rotated RNC). Source data for **Fig. 1e and Supplementary Figs. 2a-b, 5b, 5e and 9e** are available in **Supplementary Data Set 1**. All other data that support the findings of this study are available from the corresponding author upon reasonable request. The uncropped images for the westerns in the main text and supplementary figures are available in **Supplementary Data Set 2-5**.

---

**Extended Data Fig. 1.**
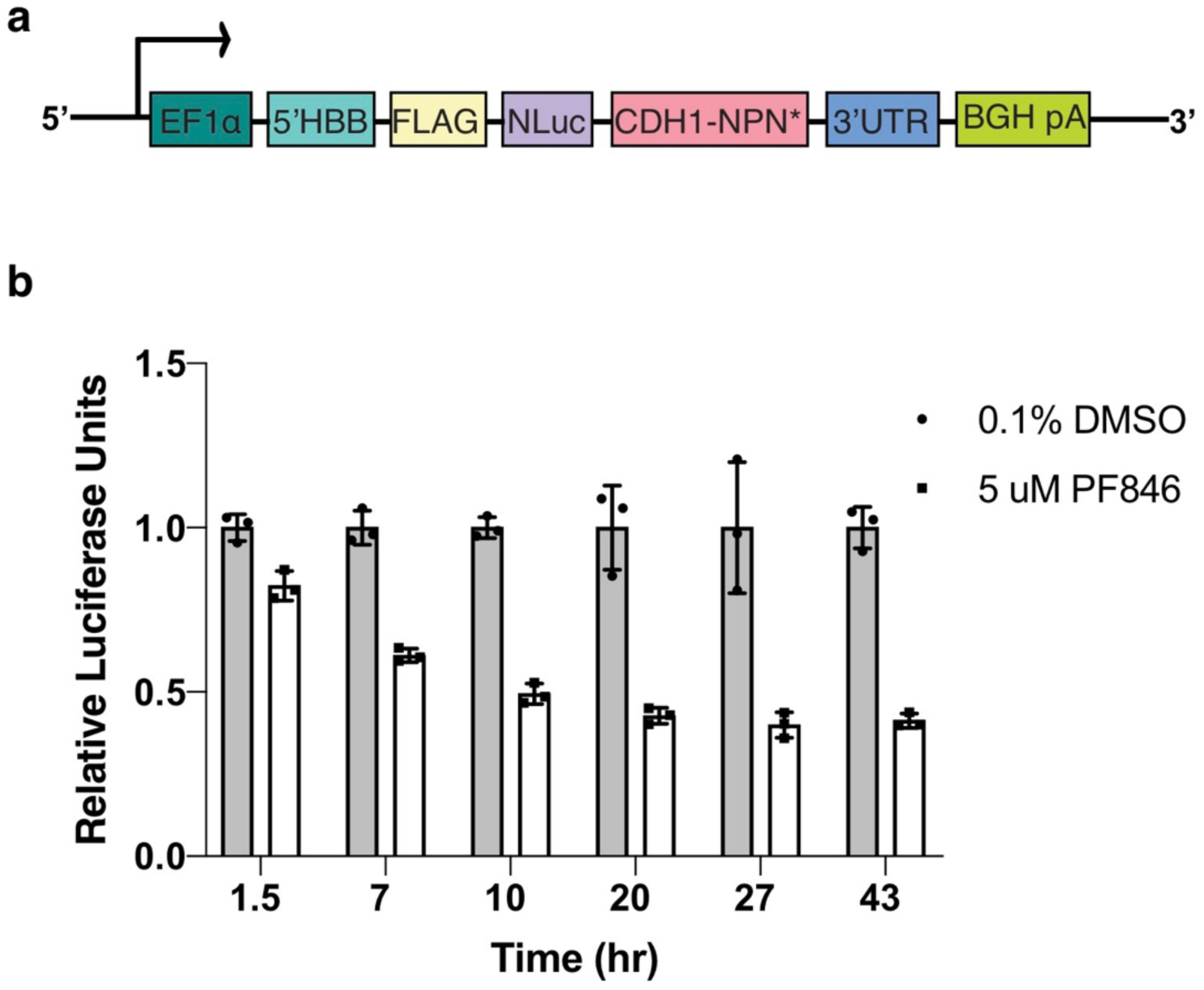
Inhibition of translation termination by PF846 in cells. (**a**) Schematic representation of the CDH1-NPN* lentiviral construct used for making stable cell lines. (**b**) Luciferase reporter assays using the CDH1-NPN* cell line at different time points after treatments with 0.1% DMSO (grey bars) or 5 μM PF846 (white bars). Bars show mean ± s.d., *n* = 3 independent experiments. Source data for (**b**) is available in **Supplementary Data Set 1**.

**Extended Data Fig. 2.**
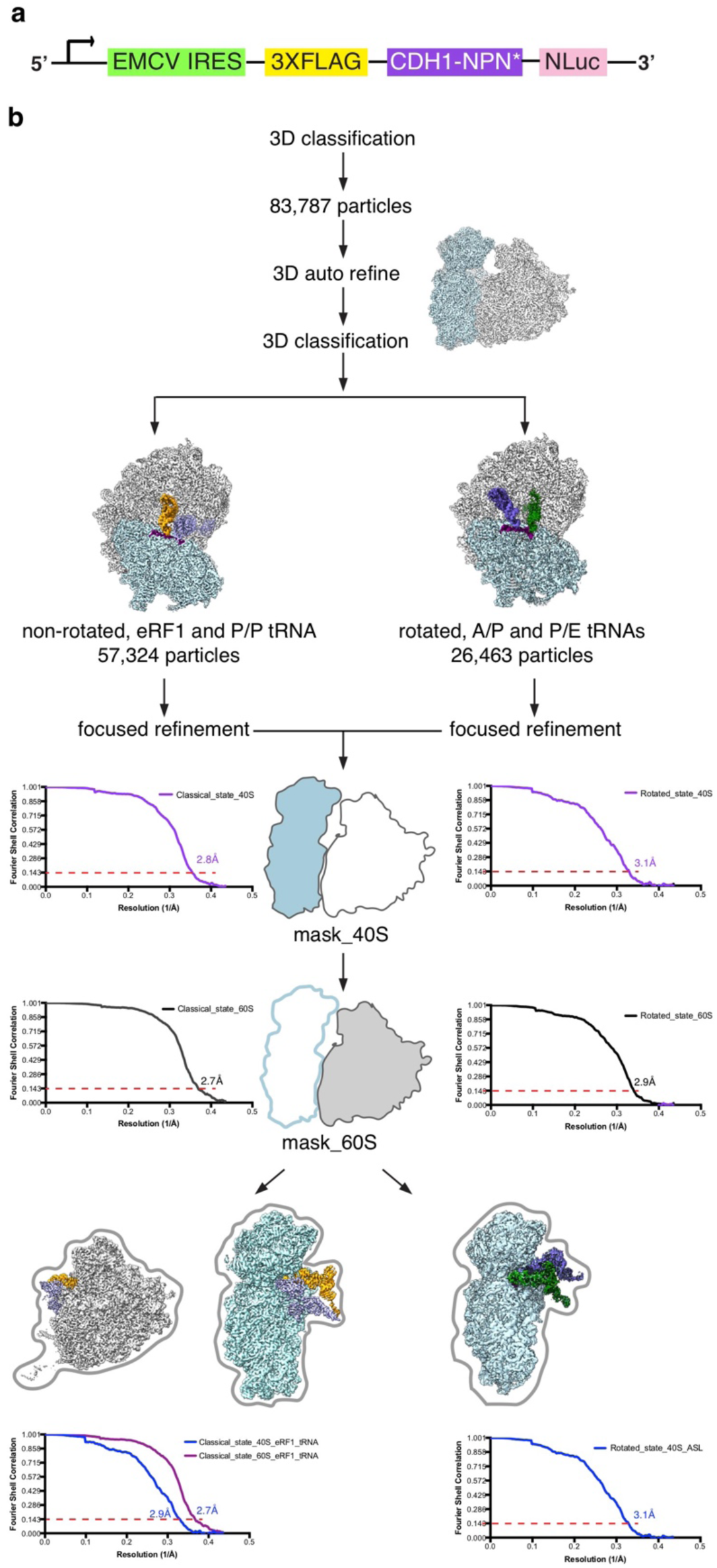
Cryo-EM data processing of PF846-stalled termination complexes. (**a**) Schematic representation of the DNA construct used to prepare PF846-stalled termination complexes. (**b**) Cryo-EM data processing workflow. Particle numbers at different steps are indicated. Two main populations were obtained after 3D classification. For the non-rotated state, focused refinements involved masking of the 40S subunit or 60S subunit individually, or the 40S subunit with eRF1, mRNA and tRNA, or the 60S subunit with eRF1 and tRNA. For the rotated state, focused refinements involved masking of the 40S subunit or 60S subunit individually, or the 40S subunit with mRNA and tRNA ASLs (anticodon stem loops). Final FSC curves of stalled RNCs in the non-rotated state and rotated state are presented, with the “gold standard” value of 0.143 used to define the resolution.

**Extended Data Fig. 3.**
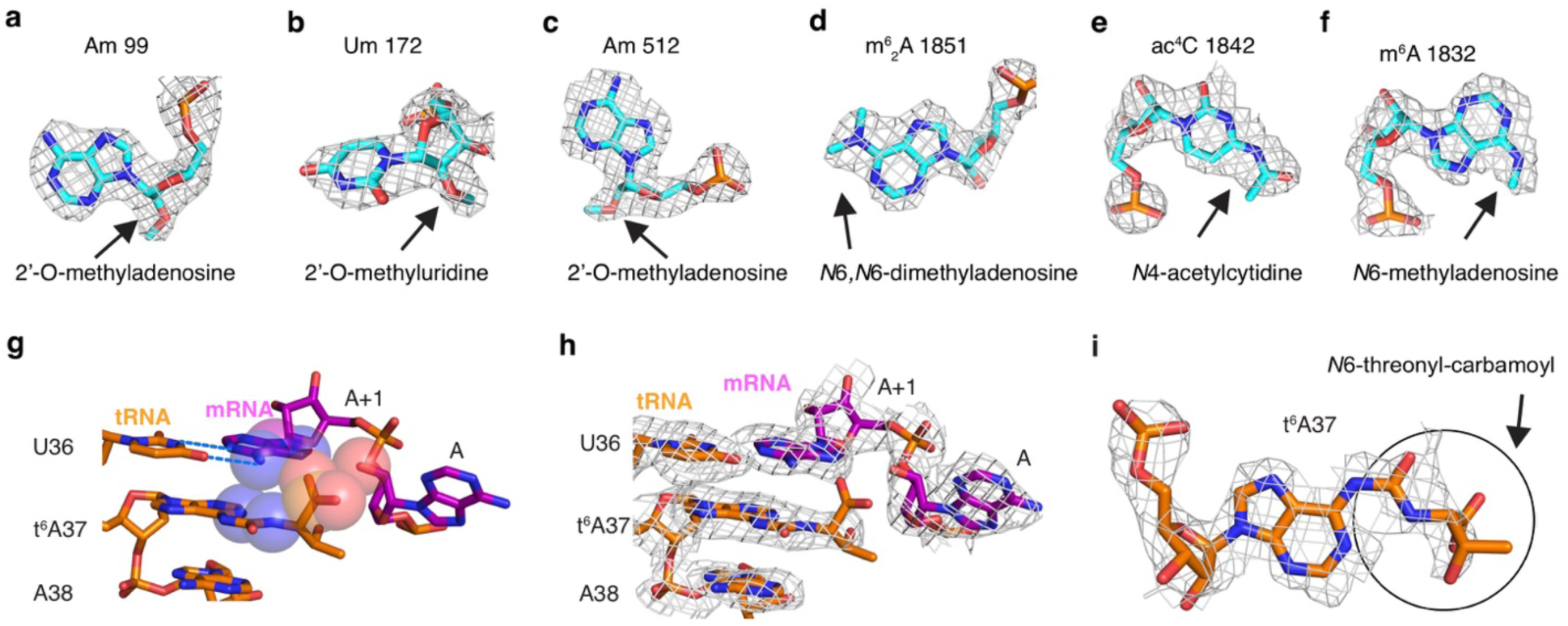
Representative rRNA and tRNA modifications from the non-rotated state. (**a-c**) Representative 18S rRNA modifications found in the non-rotated RNC, observed in mass spectrometry experiments ^15^ but not in previous cryo-EM structures. (**d-f**) Representative 18S rRNA modifications observed in this study. (**g**) Interactions between mRNA codon nucleotides A and A+1 (magenta) with P-site tRNA nucleotides 36-38 (orange). Hydrogen bonds indicated with dashed lines and van der Waals radii are shown with spheres. (**h**) Density for model shown in (**g**). (**i**) Model and cryo-EM density for *N*^6^-threonyl-carbamoyl aN6-threonyl-carbamoyl adenosine at nucleotide 37 (t^6^A37), a universal tRNA modification of almost all tRNAs decoding ANN codons (N = A,U,C or G) ^16^. The *N*-threonylcarbamoyl group is highlighted with a circle.

**Extended Data Fig. 4.**
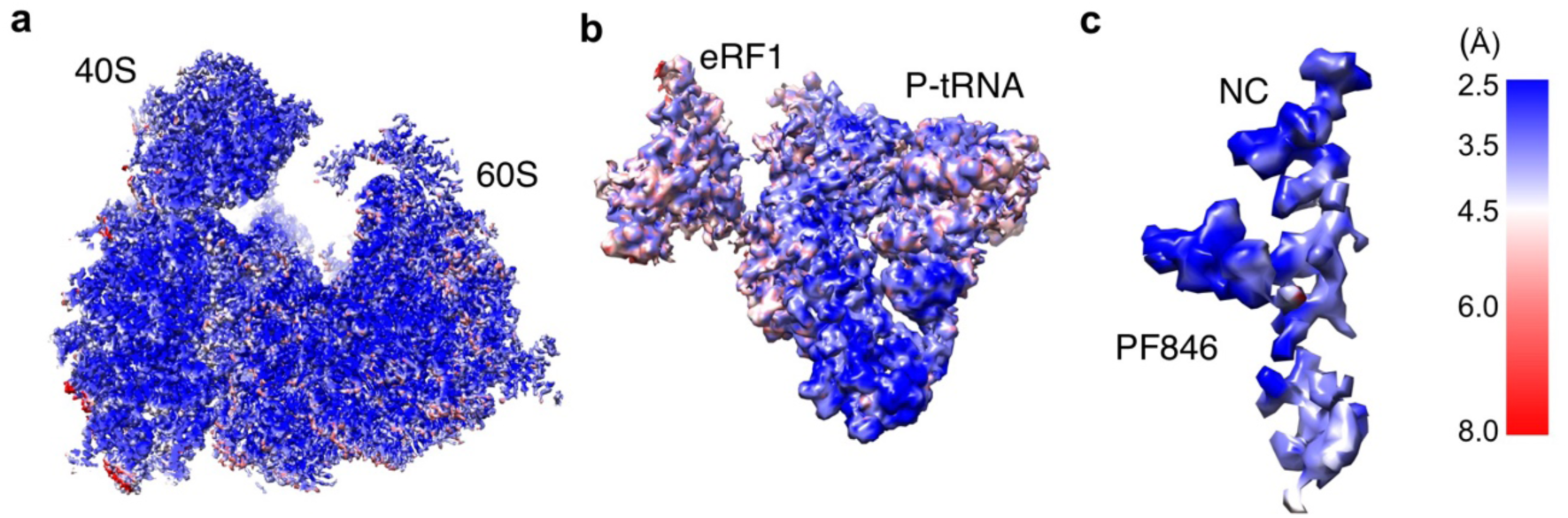
Local resolution of cryo-EM maps of PF846-stalled termination RNCs. (**a-c**) Local resolution estimation of the non-rotated RNC with (**a**) 40S subunit and 60S subunit, (**b**) 40S subunit plus tRNA plus eRF1 and (**c**) NC plus PF846. Resolution scale in angstroms (Å) is shown to the right.

**Extended Data Fig. 5.**
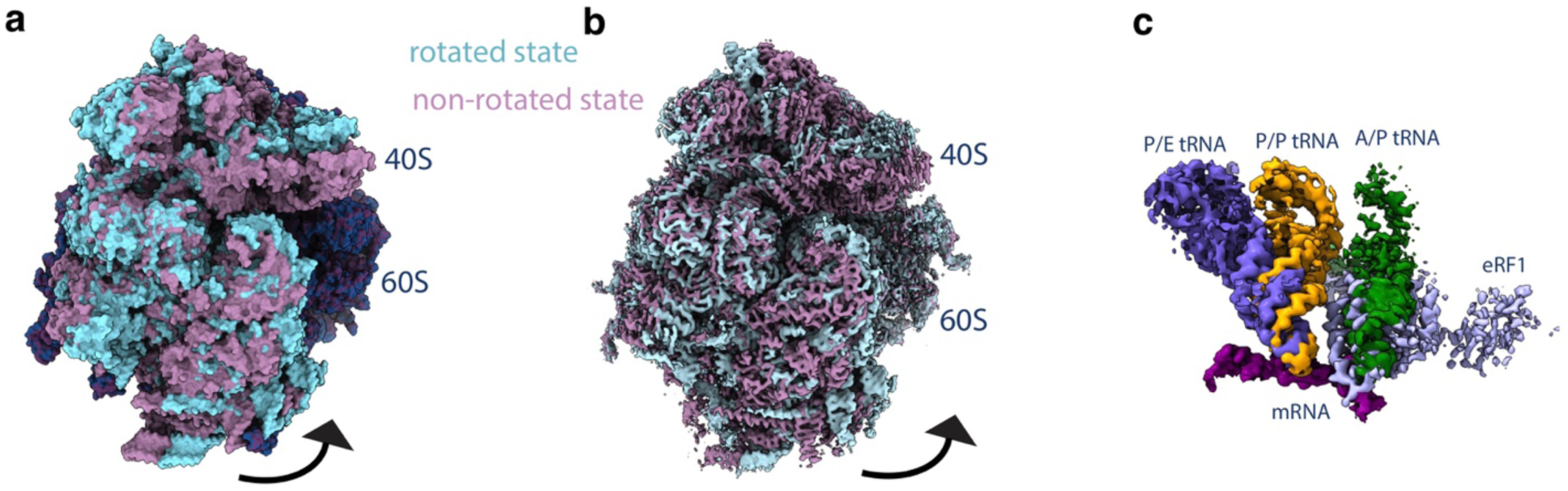
Comparison of the two different states of PF846-stalled termination RNCs. (**a**) Comparison of the atomic model of the 40S subunit in the rotated state (light cyan) with the non-rotated state (dark purple). Alignments were done using the model or map with 60S subunit as the reference. (**b**) Comparison of the cryo-EM density map of the 40S subunit in the rotated state (light cyan) with the non-rotated state (dark purple). (**c**) Comparisons of the ribosomal A, P, and E sites in both structural models, with A/P-site tRNA (dark green) and P/E-site tRNA (purple) from the rotated state; eRF1 (slate blue) and P/P-site tRNA (orange) from the non-rotated state.

**Extended Data Fig. 6.**
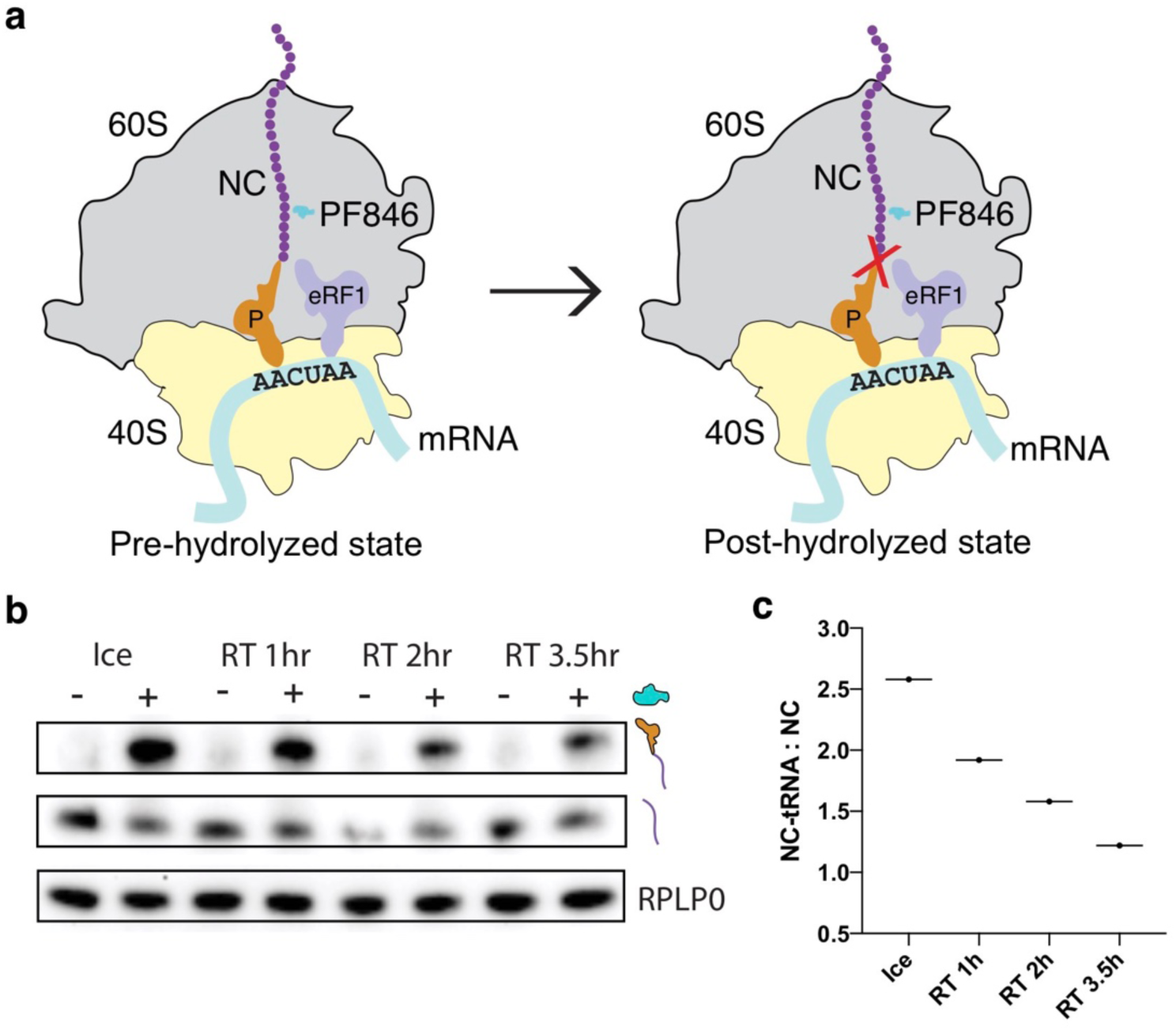
Hydrolysis of the NC-tRNA covalent bond in PF846-stalled termination complexes. (**a**) RNC models for pre- (left) and post-hydrolyzed (right) nascent chain-tRNA complexes, with hydrolysis indicated with a red “X”. (**b-c**) Western blot and quantification of NC-tRNA hydrolysis as a function of time under different conditions, ice for 3.5 hr and room temperature (RT) for 1-3.5 hr. Source data for (**c**) is available in **Supplementary Data Set 1**. Uncropped gel images for (**b**) are available in **Supplementary Data Set 3**

**Extended Data Fig. 7.**
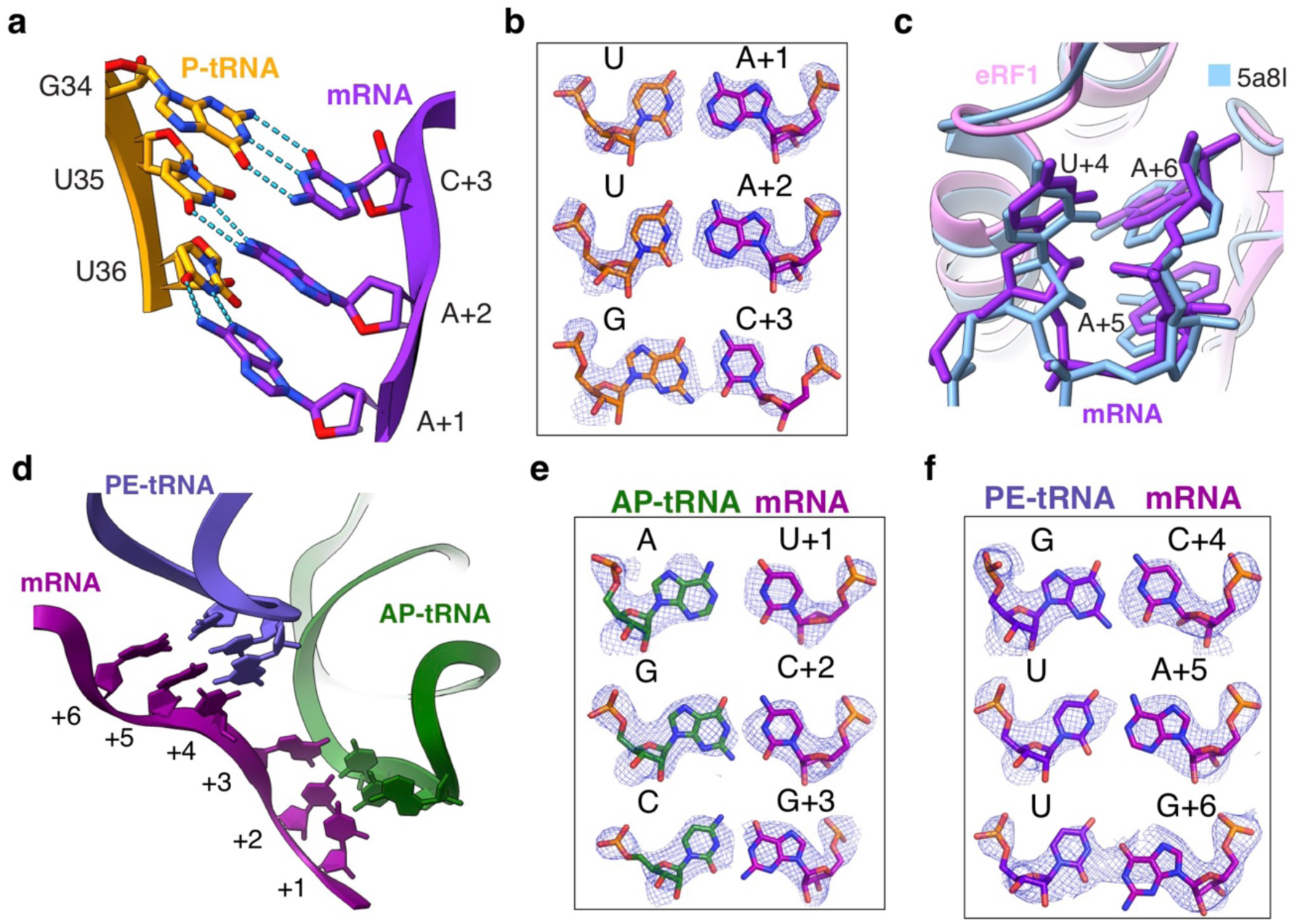
Analysis of the mRNA codon-tRNA anticodon stem loop (ASL) interactions. (**a**) Model of the P-site tRNA ASL (orange) from the non-rotated state RNC, with mRNA in magenta. The dashed lines indicate base pairing interactions. (**b**) Cryo-EM density of the P-site tRNA and mRNA in the non-rotated state, colored as in (**a**). (**c**) Superposition of the UAA stop codon with a previously-reported human termination structure (light blue) ^10^. (**d**) Model of the ASLs from the rotated RNC, with mRNA colored in magenta, P/E-site tRNA in dark purple and A/P-site tRNA in green. (**e-f**) Models of the base pairs between A/P-site tRNA anticodon nucleotides and mRNA codon nucleotides. Note that the individual nucleotides cannot be modeled with either pyrimidine or purine bases. The base pairs were refined into the cryo-EM density of the rotated RNC using Phenix.

**Extended Data Fig. 8.**
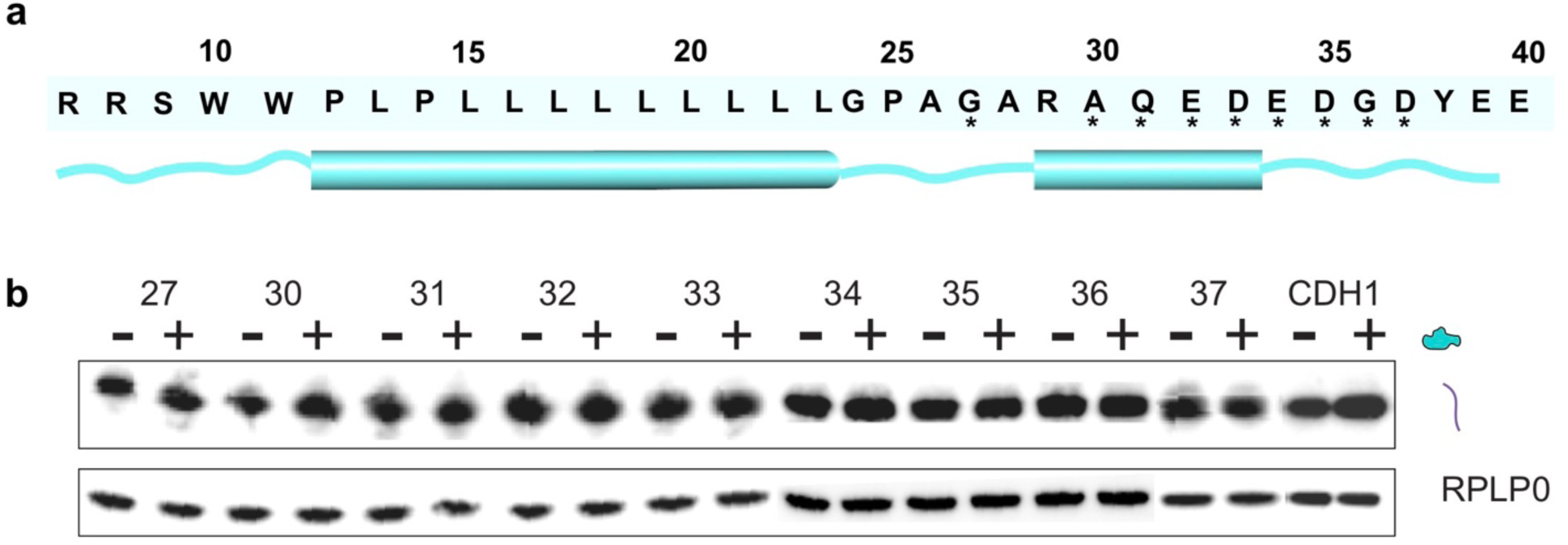
PF846-dependent PCSK9 stalling sequence with NPN* insertions. (**a**) Secondary structure prediction of the PCSK9 nascent chain sequence required for PF846-dependent stalling of elongation, using Phyre2 ^61^, with α-helices represented by cylinders and loops as curved lines. The stalling sequence of PCSK9 is shown. Asterisks indicate the beginning amino acid for NPN* replacements used in translation assays. (**b**) Western blot showing the effects of NPN* insertions at different positions in the PCSK9 nascent chain. *In vitro* translation reactions with DMSO (–) or with 50 μM PF846 (+) are shown. Samples were treated with RNase A, and the stalled PCSK9 nascent chains were detected by blotting with an anti-FLAG antibody, with RPLP0 serving as a loading control. Uncropped gel images for (**b**) are available in **Supplementary Data Set 4**.

**Extended Data Fig. 9.**
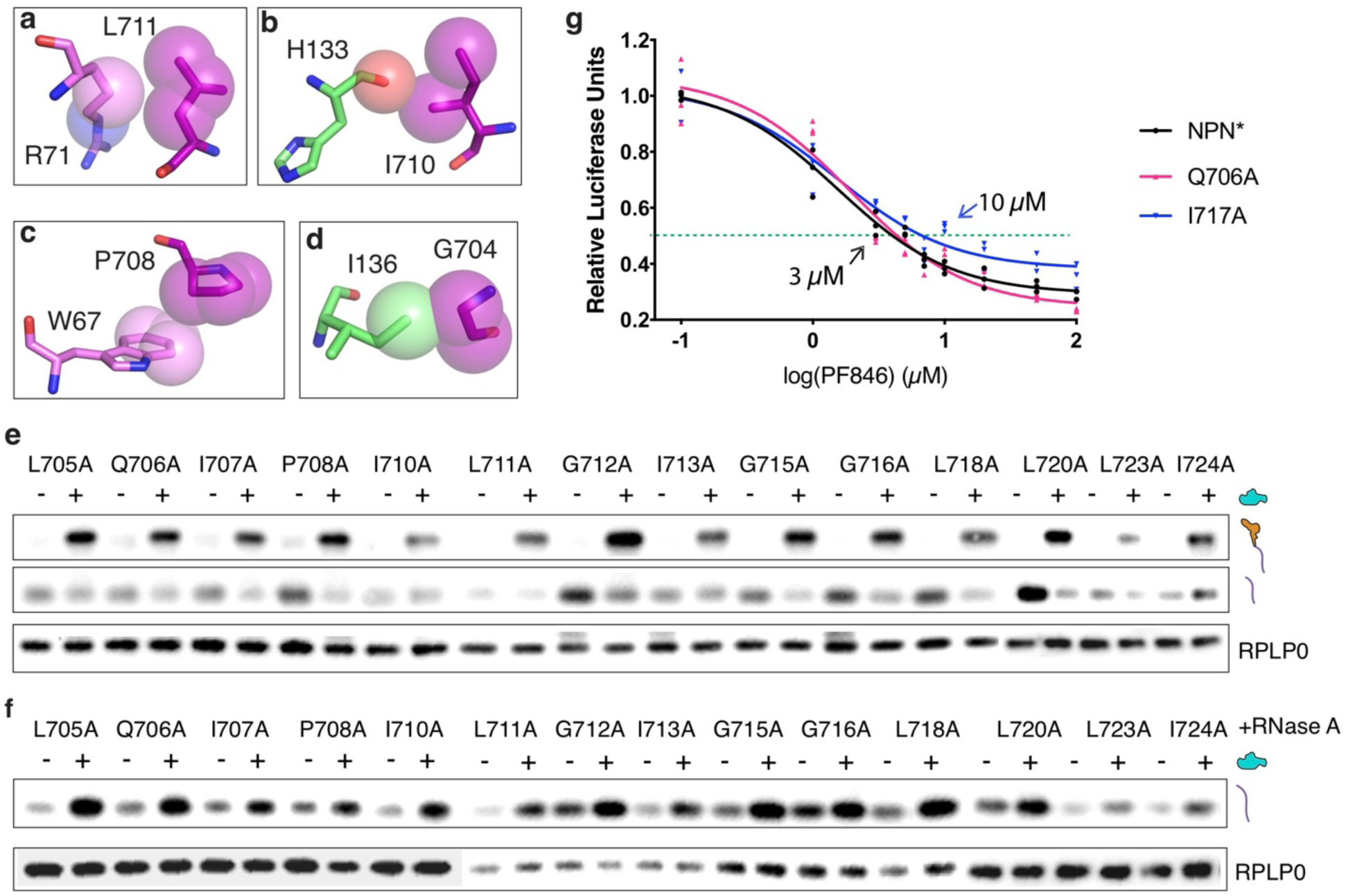
Effects of mutations in the CDH1-NPN nascent chain on PF846-dependent inhibition of termination. (**a-d**) Interactions of ribosomal protein uL22 (light green) and uL4 (hot pink) with the nascent chain, with van der Waals surfaces shown. (**e**) Western blots of FLAG-tagged CDH1-NPN nascent chains containing single mutations, from *in vitro* translation reactions in the presence (+) or absence (–) of 50 μM PF846. The positions of tRNA-bound and free nascent chains are shown, with RPLP0 serving as a loading control. RNCs were assembled as in **Figure 3e**. (**f**) Western blots with RNC samples from (**e**) after treatment with RNase A. (**g**) IC50 values for PF846-dependent inhibition of translation termination, using stable cell lines expressing the CDH1-NPN* nascent chain (black dots) or NCs with mutations Q706A (red dots) and I717A (blue dots). Source data for (**g**) are available in **Supplementary Data Set 1**. Uncropped gel images for (**e**) and (**f**) are available in **Supplementary Data Set 5**.

**Extended Data Fig. 10.**
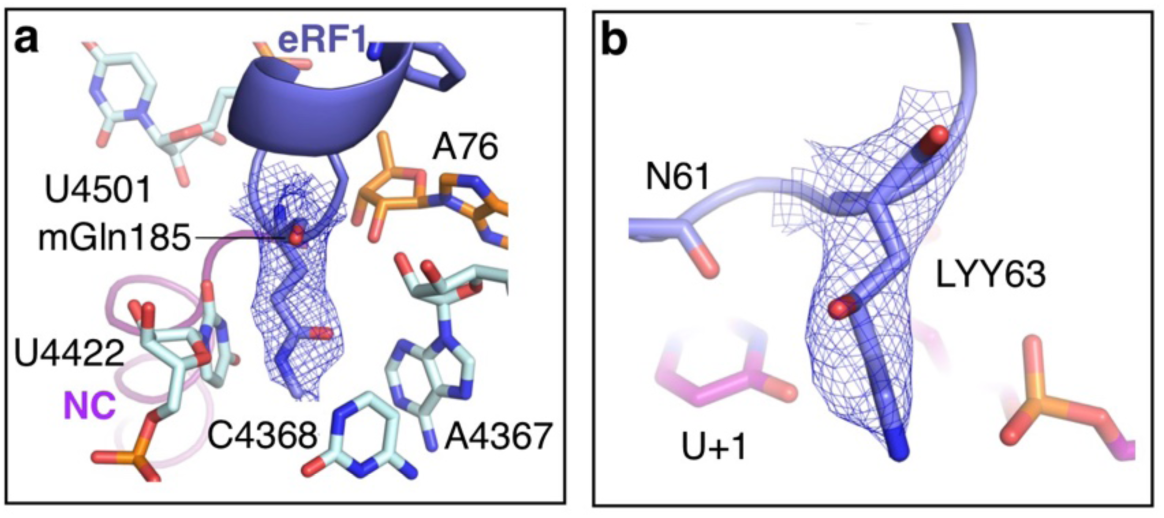
Posttranslational modifications of eRF1 observed in the cryo-EM density map. (**a**) Close-up view of the GGQ motif within the PTC. Cryo-EM density for the methylated Gln (mGln185, slate blue) of the GGQ motif, positioned next to A76 of P-site tRNA and surrounded by PTC rRNAs, is represented with mesh. The map was sharpened with a B-factor of -40 Å^2^. (**b**) The cryo-EM density for the C4 hydroxylysine 63 (LYY63) shown in mesh. The map was sharpened with a B-factor of - 40 Å^2^.

**Extended Data Fig. 11.**
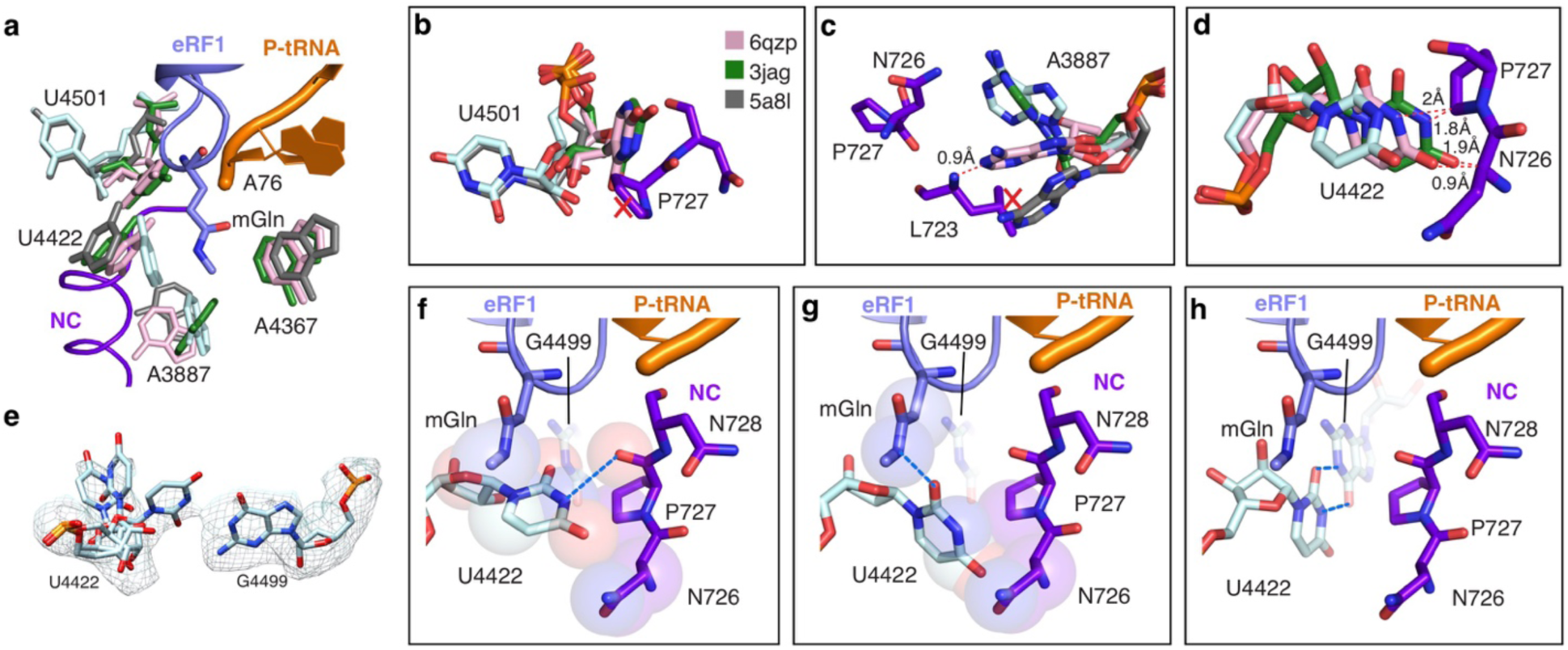
Conformation of the PTC in eukaryotic ribosome termination complexes. (**a**) Overall alignment of the PTC from different eukaryotic ribosome structures (6qzp ^14^ in pink, 3jag ^9^ in forest green, and 5a8l ^10^ in grey). Positions of the eRF1 GGQ motif (slate blue), the P-site tRNA (orange) and nascent chain (purple) are shown. (**b-d**) Positions of (**b**) U4501, (**c**) A3887 and (**d**) U4422 aligned within the PTC. Potential steric clashes are highlighted with a red “X” or with labeled atomic distances. (**e**) Cryo-EM density for U4422 and G4499, with U4422 modeled in three different conformations based on the observed density. (**f**) The major conformation of U4422 observed in the cryo-EM density, which makes multiple interactions with the nascent chain and mGln185 of eRF1. Dashed lines indicate hydrogen bonds and spheres represent van der Waals radii. (**g**) Conformation showing U4422 turned outward from the nascent chain, with H-bonds and van der Waals radii as in (**g**). (**h**) Conformation showing U4422 flipped inward to base pair with G4499.

**Extended Data Fig. 12.**
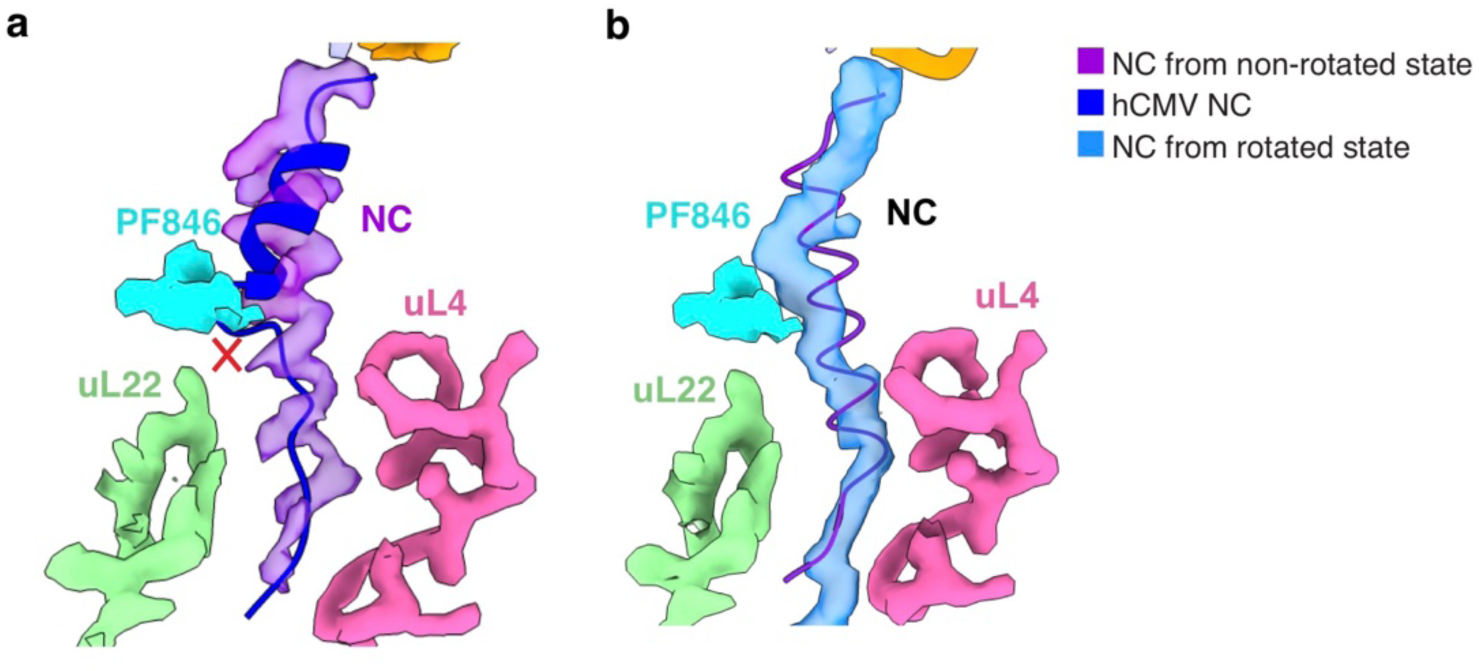
Conformation of NCs within ribosomes stalled at termination. (**a**) Superposition of the hCMV nascent chain model (blue, PDB: 5a8l) ^10^ with the ribosome exit tunnel of the non-rotated CDH1-NPN RNC stalled by PF846. The potential steric clash of the hCMV NC with PF846 is indicated. (**b**) Superposition of the CDH1-NPN NC model from the non-rotated RNC with the cryo-EM density of the rotated RNC.

**Extended Data Table 1 | DNA primers used in this study.**

**Supplementary data set 1 | Source data for the bar graphs presented in the main text.**

**Supplementary data set 2 | The uncropped images for gels shown in Fig.3e.**

**Supplementary data set 3 | The uncropped images for gels shown in Extended Data Fig. 6b.**

**Supplementary data set 4 | The uncropped images for gels shown in Extended Data Fig. 8b.**

**Supplementary data set 5 | The uncropped images for gels shown in Extended Data Fig. 9e-f.**

