## Supplementary Data Sets 2-5 for "Selective inhibition of human translation by a drug-like compound that traps terminated protein nascent chains on the ribosome"

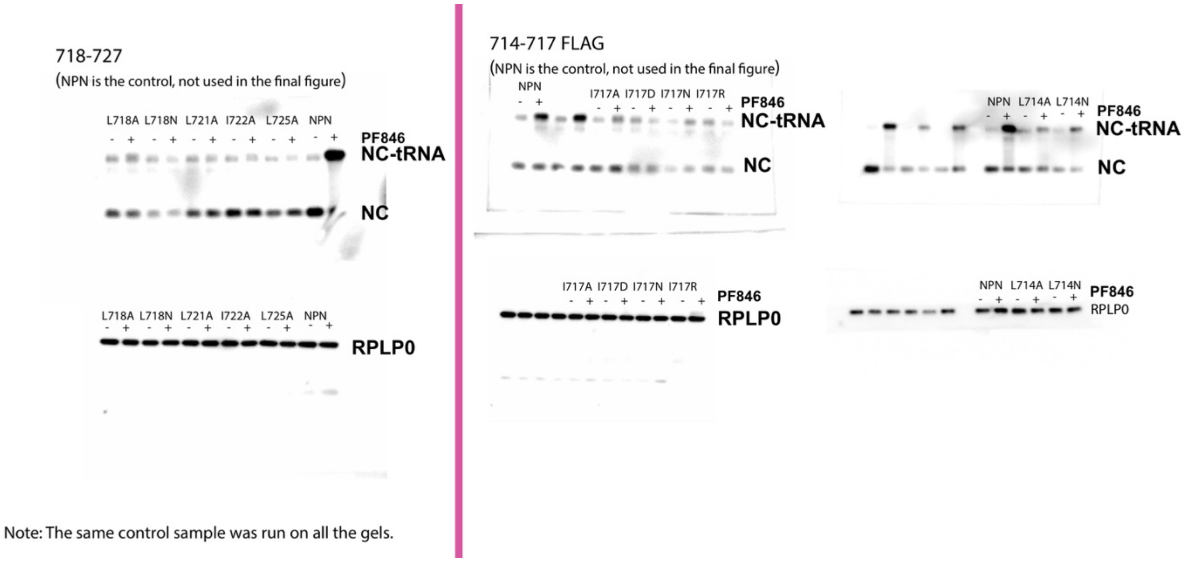


**Supplementary data set 2 | The uncropped images for gels shown in Fig.3e.**

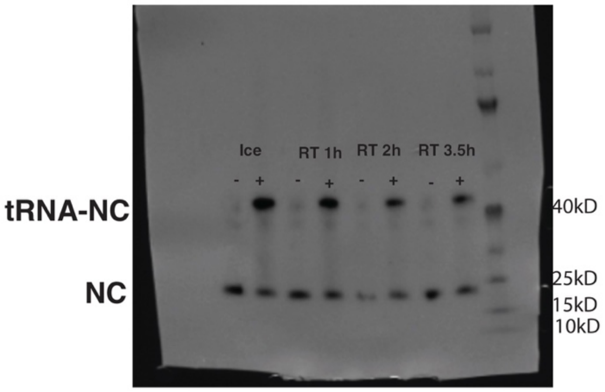


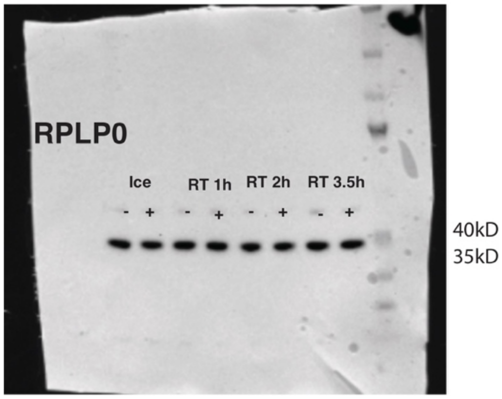


**Supplementary data set 3 |** The uncropped images for gels shown in **Extended Data Fig. 6b.**

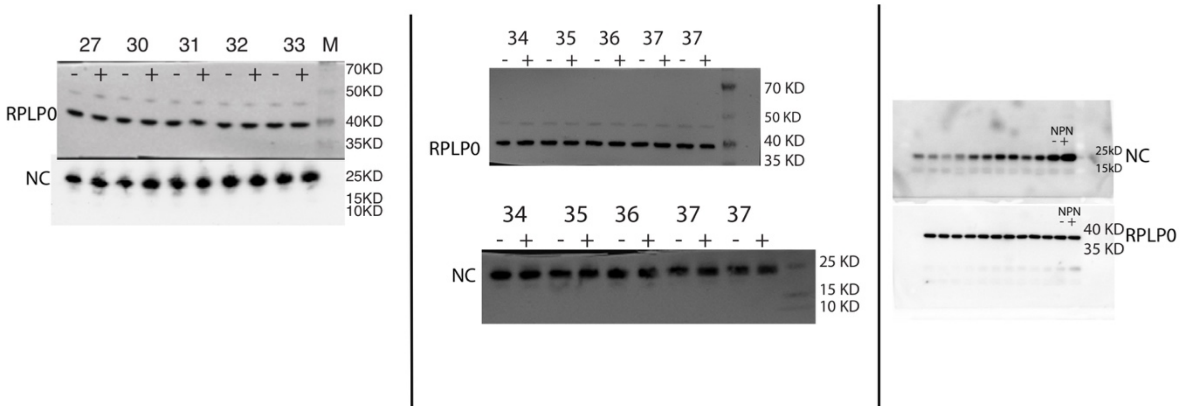


**Supplementary data set 4 | The uncropped images for gels shown in Extended Data Fig. 8b.**

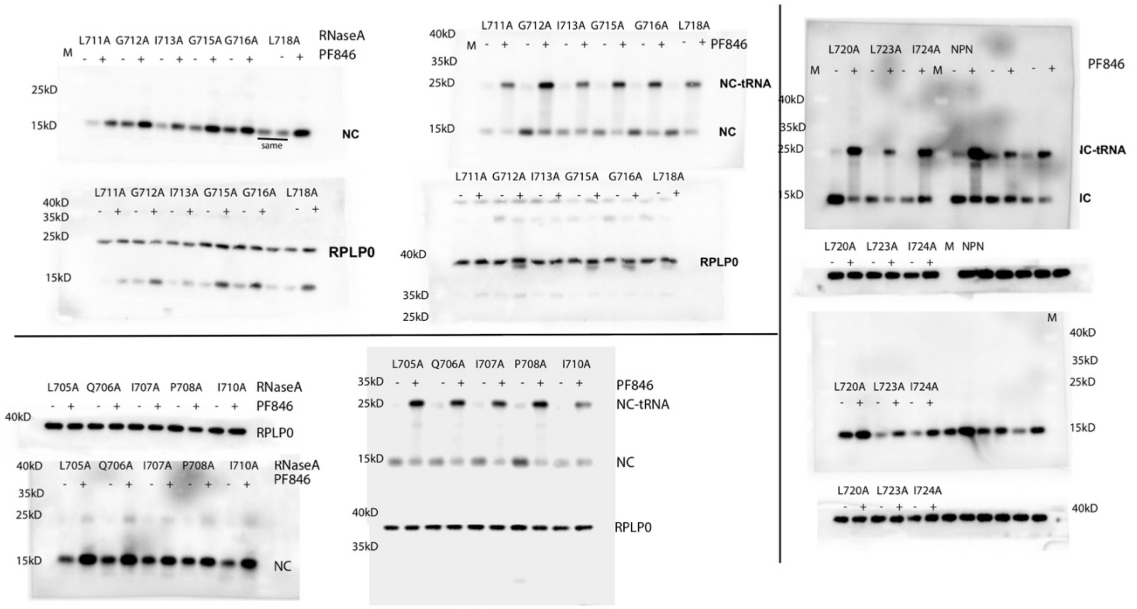


**Supplementary data set 5 | The uncropped images for gels shown in Extended Data Fig. 9e-f.**
